# Nanoscale remodeling of synaptic glutamate receptors and calcium channels in temporal lobe epilepsy

**DOI:** 10.64898/2026.10.21.760440

**Authors:** Walter A. Kaufmann, Alessandro Venturino, David Kleindienst, Karl Roessler, Thomas Czech, Jan Maximilian Janssen, Maren Engelhardt, Harald Stefanits, Masahiko Watanabe, Sandra Siegert, Ryuichi Shigemoto

**Author notes:** Correspondence to: Walter Anton Kaufmann, Institute of Science and Technology Austria (ISTA), A-3400 Klosterneuburg, Am Campus 1, Austria. **Senior authorship:** Sandra Siegert and Ryuichi Shigemoto share senior authorship.

## Abstract

Temporal lobe epilepsy (TLE) is the most common form of focal epilepsy and remains drug-resistant in approximately one-third of patients, for whom surgical resection of the epileptogenic zone is currently the most effective treatment option. One strategy to improve therapeutic outcomes involves identifying the molecular mechanisms and key regulators underlying synaptic function. Although altered glutamatergic neurotransmission is considered a major contributor to epileptic hyperexcitability, the nanoscale organization of key synaptic proteins in the human epileptic cortex is still not well characterized. In particular, it remains unclear whether pre- and postsynaptic proteins undergo synapse-type-specific remodeling in TLE.

Here, we used high-resolution freeze-fracture replica immunolabeling to map the nanoscale organization of postsynaptic AMPA receptors (AMPARs) and presynaptic P/Q-type voltage-gated calcium channels (Ca_V_2.1) in glutamatergic synapses of cortical layers II/III of surgically resected human tissue. We examined perilesional, cytologically unremarkable, non-epileptogenic cortical tissue obtained from patients undergoing tumor resection (52-63 years; two female and three male patients) and compared these data with cortical samples from patients with refractory TLE (22-53 years; two female and four male patients). To assess whether pathological remodeling differs across synapse populations, we analyzed both excitatory dendritic spine and shaft synapses.

We found a robust, highly consistent increase in AMPAR density at both spine and shaft synapses in TLE patients indicating enhanced postsynaptic excitatory signaling. However, AMPAR spatial reorganization differed between synapse types. Monte Carlo simulations demonstrated that spine synapses largely preserved a random receptor distribution without a significant clustering pattern, whereas shaft synapses exhibited a redistribution from clustering toward random receptor arrangements together with reduced center-preference within the postsynaptic density.

In contrast, presynaptic Ca_V_2.1 remodeling was more variable and predominantly associated with excitatory spine synapses. TLE spine synapses exhibited an overall increase in Ca_V_2.1 density together with significantly larger and more numerous Ca_V_2.1 nanoclusters within the active zone. This difference was particularly pronounced in male patients. In comparison, shaft synapses displayed only minor presynaptic alterations. Ca_V_2.1 channels remained highly clustered under both control and epileptic conditions, indicating preservation of core active-zone organizational features despite disease-associated nanoscale remodeling.

Together, these findings provide direct ultrastructural support that pathological plasticity in human TLE involves an overall increase in excitatory postsynaptic responsiveness and a target cell-specific increase of excitatory release probability at cortical synapses.

## Introduction

Epilepsy is one of the most common chronic neurological disorders, affecting approximately 50 million people worldwide.^1^ Although antiseizure medications provide symptomatic seizure control for many patients, approximately one-third develop drug-resistant epilepsy.^2^ The marked heterogeneity in epilepsy epidemiology, etiology, pathophysiology, and treatment responses further complicates clinical management.^3^ For appropriately selected patients with drug-resistant focal epilepsy, resective microsurgery remains the most effective treatment option and is associated with substantially higher seizure-freedom rates than continued pharmacotherapy alone. Long-term cohort studies and meta-analyses indicate that surgery results in seizure freedom in approximately 60-70% of selected patients.^4–6^ Nevertheless, many patients are ineligible for surgery or do not achieve sustained seizure freedom, underscoring the need to identify alternative pharmacological targets and strategies through a deeper understanding of the synaptic mechanisms that contribute to seizure generation and network dysfunction in temporal lobe epilepsy (TLE).

Glutamatergic neurotransmission plays a central role in the initiation and propagation of seizures and in the long-term pathophysiology of epilepsy.^7,8^ Excessive excitation and disruption of the excitation-inhibition balance contribute to recurrent seizures and maladaptive alterations in synaptic structure and function, and network connectivity.^9,10^ These pathological processes involve coordinated remodeling of both presynaptic release mechanisms and postsynaptic glutamate receptor organization at excitatory synapses.^8^ Among ionotropic glutamate receptors, the α-amino-3-hydroxy-5-methyl-4-isoxazolepropionate (AMPA) receptors (AMPAR) mediate the majority of fast excitatory neurotransmission in the brain^11,12^ and are major determinants of synaptic strength and excitability. These receptors are tetrameric complexes in homomeric or heteromeric forms composed of the four subunits GluA1-GluA4.^13^ Accordingly, AMPARs are a clinically validated target for the treatment of focal-onset seizures, including seizures associated with TLE.^14,15^

In parallel, presynaptic voltage-gated calcium channels (VGCCs) critically regulate glutamatergic transmission, particularly Ca_V_2.1-containing P/Q-type channels, which control Ca^2+^-dependent synaptic vesicle release.^16,17^ They are the principal source of Ca^2+^ influx required for neurotransmitter release at many central synapses and play a predominant role in the neocortex.^18^ Moreover, mutations in the gene encoding the Ca_V_2.1 α1 subunit (*CACNA1A*) have been implicated in various forms of seizures and status epilepticus, and several anti-seizure medications act, at least in part, by modulating Ca_V_2.1 channels.^19^ Together, dysregulation of both AMPARs and Ca_V_2.1 channels has been critically implicated in epilepsy and may contribute to pathological synaptic plasticity and network hyperexcitability.

Synaptic transmission is essentially determined not only by the abundance of synaptic proteins, but also by their precise nanoscale organization within pre- and postsynaptic membrane specialization. Excitatory cortical synapses display substantial structural and molecular heterogeneity, including distinct spine and shaft synapse populations that differ in their functional properties and may undergo differential pathological remodeling.^20^ Although altered glutamatergic signaling has long been implicated in epilepsy, the nanoscale organization of key synaptic proteins in human epileptic synapses remains poorly understood. In particular, it is unknown whether postsynaptic AMPARs and presynaptic Ca_V_2.1 channels undergo synapse-type-specific reorganization in TLE.

To address these questions, we applied high-resolution freeze-fracture replica immunolabeling to analyze the nanoscale distribution of AMPAR and Ca_V_2.1 channels in glutamatergic synapses of cortical layer II/III from patients with TLE. We found distinct forms of pre- and postsynaptic nanoscale remodeling at excitatory synapses, including increased AMPAR density in both spine and shaft synapses, subtle synapse-type-dependent differences in AMPAR spatial organization, and selective upregulation and enhanced clustering of Ca_V_2.1 channels in spine synapses. Together, these findings reveal synapse type-specific remodeling of excitatory synapses in human TLE and provide new insights into the synaptic mechanisms underlying cortical hyperexcitability in epilepsy.

## Materials and methods

### Subject details

This study includes samples from nine individuals admitted to neurosurgery at the Medical University of Vienna, Department of Neurosurgery (Vienna, Austria) and two individuals admitted to neurosurgery at the Kepler University Hospital Linz, Department of Neurosurgery (Linz, Austria). All patients underwent a comprehensive clinical evaluation before surgery, and the resections were performed strictly for therapeutic reasons. The patients’ consent was obtained for the use of brain surgical specimens and clinical data for research purposes according to the Declaration of Helsinki. Experimental procedures were evaluated and certified by the Ethics Committee of the Medical University of Vienna (authorizations EK 1188/2019 and EK 2271/2021) and Johannes Kepler University Linz (authorization EK 1250/2023).

We first examined perilesional, cytologically unremarkable, non-epileptogenic cortical tissue from five patients undergoing resection for brain tumors, ranging in age from 52 to 63 years (mean age: 58 years), including two females and three males (hereafter referred to as controls). We then analyzed cortical tissue samples from six patients with drug-resistant TLE, ranging in age from 22 to 53 years (mean age: 39 years), including two females and four males. Clinical characteristics and histopathological diagnosis are summarized in Table 1. The control cohort included patients with brain metastases, oligodendroglioma, or other gliomas; histopathological assessment found no evidence of tumor infiltration in the sampled cortical tissue.

**Table 1:** Patient list (pseudonymized data)

| Pt # | ID | Sex | Age | Diagnosis | Histology |
| --- | --- | --- | --- | --- | --- |
| ***Non-epileptogenic control patients*** |  |  |  |  |  |
| I | Pt7 | F | 52 | Brain metastasis | Cerebral cortex and subcortical white matter with fibrinoid necrosis, vascular occlusion, and vessel-wall necrosis, consistent with therapy-associated changes; no residual adenocarcinoma identified |
| II | Pt18 | M | 59 | Oligodendro-glioma WHO Grade II | Mild meningeal fibrosis with focal reactive arachnoid-cell proliferation; no tumor infiltration |
| III | Pt26 | M | 58 | Glioma | Preserved cortical layering and architecture with marked subpial fibrous gliosis and focal meningeal thickening; no abnormal SMI-31, SMI-32, or CD34 staining and no vascular proliferation |
| IV | Pt31 | F | 56 | Low Grade Glioma | Surgical access-path tissue with preserved cellular architecture; GFAP and S100 positively without increased Olig2 expression; no tumor infiltration |
| V | Pt32 | M | 63 | Glioma WHO Grade I | Cortical access-path tissue without vascular proliferation or tumor infiltration |
| ***Temporal lobe epilepsy patients*** |  |  |  |  |  |
| VI | Pt3 | F | 35 | TLE | No histopathological abnormalities identified |
| VII | Pt4 | M | 53 | TLE | No histopathological abnormalities identified |
| VIII | Pt6 | M | 52 | TLE | Mild diffuse gliosis, most pronounced subpially, with preserved cortical layering and partially enlarged perivascular spaces; no definite lesion identified |
| IX | Pt16 | M | 22 | TLE | Mild meningeal fibrosis with sparse lymphocytic infiltration; mild oligodendroglial reactivity in the subcortical white matter and a mildly blurred corticomedullary junction with increased ectopic neurons in subcortical layers; otherwise unremarkable |
| X | Pt17 | F | 31 | TLE | Mild ribbon-like subpial fibrous gliosis; otherwise unremarkable |
| XI | Pt25 | M | 42 | TLE | Minimal gliosis in the superficial cortical layers; otherwise unremarkable |

### Excision of samples: temporal cortex

In non-epileptic patients undergoing neurosurgery for removal of a temporal tumor and in TLE patients undergoing neurosurgery for resection of the epileptogenic zone, a cortical sample was excised during the surgical approach to the underlying diseased brain tissue. After sharp dissection of the pial surface with a scalpel and micro-scissors, an approximately 1-cm^3^ tissue block comprising cortex and subcortical white matter was removed with a blunt dissector without cauterizing the tissue.

### Fixation of surgery samples

Immediately after resection, and within no more than 10 minutes, the excised tissue samples were immersed in physiological saline solution (0.9% NaCl (w/v) in water), brought to the laboratory, and fixed within 15 min after excision by immersion in a 2% formalin solution (2% paraformaldehyde (w/v) and 15% of saturated picric acid solution (v/v) in 0.1 M phosphate buffer (PB); pH 7.4) overnight at 4 °C. The samples were then stored in 0.1 M PB containing 0.02% sodium azide (NaN_3_; w/v) at 4 °C until further processing.

### SDS-digested freeze-fracture replica labeling (SDS-FRL)

For the high-resolution localization of the proteins of interest, we applied sodium dodecyl sulfate (SDS)-digested freeze-fracture replica labeling (FRL) electron microscopy with some modifications to the original method.^21^ Freeze-fracture replica preparation causes membrane splitting along the hydrophobic core of the lipid bilayer, where the physical tension is lowest. This process exposes the inner, protoplasmic (P-face) and the outer, extracellular (E-face) membrane leaflets, thereby allowing replicas of these surfaces to be prepared for immunolabeling and analysis at the transmission electron microscopy (TEM) level.^22^

Slices perpendicular to the cortical surface were cut at 140 µm with a VT1200 vibroslicer (Leica Microsystems) and cryo-protected by immersion in 30% glycerol in PB (v/v) overnight at 4 °C. The slices containing the cortical layer II/III were trimmed into 1.2 mm diameter circles, sandwiched between copper carriers, and rapidly frozen by a high-pressure freezing machine HPM 010 (Leica Microsystems). The slices were then fractured by a double-replica method as described before for rodent tissue samples.^22,23^ The frozen slices were mounted on a double-replica table for tensile fracture (Leica Microsystems) under liquid nitrogen and inserted into the freeze-etching device EM ACE900 (Leica Microsystems). The slices were fractured at a temperature of -120 °C and chamber pressure of 8 x 10E-7 mbar. Fractured faces were first replicated by evaporation of carbon (rotating) by means of an electron beam gun positioned at a 90° angle to a thickness of 5 nm for the immobilization of macromolecules. The faces were then shadowed unidirectionally with platinum-carbon at a 60° angle (thickness 2 nm) and strengthened by evaporation of carbon to a thickness of 20 nm from a 90° angle (rotating). Samples were brought to atmospheric pressure and thawed to room temperature (RT) in Tris-buffered saline (TBS; 50 mM, 0.9% NaCl (w/v); pH 7.4). Tissue not immobilized in the replica membrane was solubilized in a medium containing 2.5% SDS (w/v) and 20% sucrose (w/v) in 15 mM TBS, pH 8.3, on a shaking platform (50 rpm) for 60 h at 60 °C. Replicas were brought to RT and kept in the same solution until further use.

For immunolabeling, the SDS solution was gradually replaced with 5% polyethylene glycol (PEG; MW 2000; w/v) in TBS, as previously described.^22^ Replicas were then incubated in 5% bovine serum albumin (BSA; w/v) in TBS for 90 min at RT to block nonspecific binding. Subsequently, primary antibodies (as specified in Supplementary Table 1) and gold-conjugated secondary antibodies were applied sequentially in TBS containing 2% BSA (w/v), incubated overnight at 15 °C on a rocking platform. The first set of primary antibodies applied was: guinea pig polyclonal anti-Ca_V_2.1 subunit of the P/Q-type voltage-gated calcium channel (Synaptic Systems), 2 µg/ml; guinea pig polyclonal anti-panAMPA receptor (Frontier Institute co.,ltd), 4 µg/ml. The antibody was generated using amino acid residues 717-754 and affinity purified with amino acid residues 727-745 [EQRKPCDTMKVGGNLDSKG] of mouse GluA1, which is common for all of GluA1-4.^24^ Replicas were then reacted with goat anti-guinea pig secondary antibodies conjugated to 5-nm gold particles (BBI) at a dilution of 1:30. Replicas were then incubated with the second primary antibody, rabbit anti-vGLUT1 (Synaptic Systems), at a concentration of 4 µg/ml. They were subsequently reacted with goat anti-rabbit secondary antibodies conjugated to 15-nm gold particles (BBI) at a dilution of 1:30. The specificity of the Ca_V_2.1 was confirmed by testing cerebellar tissue samples from Ca_V_2.1 knock-out mice.^25^ The specificity of secondary antibodies was tested by omitting primary antibodies and applying the full set of secondary antibodies.

Replicas were then washed in water, mounted onto Formvar-coated 100-line copper grids (Electron Microscopy Sciences), and examined in a Tecnai 12 TEM (Thermo Fisher Scientific) operated at 120 kV and equipped with a side-mounted Veleta CCD camera (EMSIS). Images were acquired using Radius (EMSIS), and analyzed using FIJI (distributed under the General Public License, GPL) and MatLab software (MathWorks).

### Sampling

One replica per patient was included in each immunolabeling experiment, and six experimental runs were performed for each protein of interest (n = 6 replicas per patient). Ten aspects of cortical layers II/III were randomly selected in each replica at a magnification of x420, corresponding to an area of 25,000 µm^2^ (total sampling area per replica = 250,000 µm^2^). This yielded a total sampling area of 1,500,000 µm^2^ per patient and protein of interest. Within each sampling area, profiles of pyramidal neurons and cortical interneurons were identified based on morphological criteria,^26^ and all excitatory synapses onto dendritic spines of pyramidal neurons and dendritic shafts of interneuron were sampled. Images of the postsynaptic membrane specialization (postsynaptic density, PSD) on the E-face and the presynaptic active zone (AZ) on the P-face were acquired at magnifications ranging from x46k to x97k. The tilt angle was adjusted for each synapse to maximize area. For the analysis of Ca_V_2.1 channels, only complete synapses were included, whereas for the AMPA receptor analysis, both complete and incomplete synapses were evaluated, because labeling densities and distribution patterns did not differ significantly between these categories. For each patient, across all replicas, we collected 46-96 PSD profiles from spine synapses, 41-60 PSD profiles from shaft synapses, 52-95 AZ profiles from spine synapses, and 43-69 AZ profiles from shaft synapses.

### Analysis

For a minimally biased and reproducible analysis of immunolabeling, Darea – a deep learning-assisted replica analysis software^27^ – was used to accelerate and standardize the analysis.^22^ Since this deep neural network was originally trained on mouse neuron synapses, we retrained the system using images of human cortical synapses, as described in Figure 2. Characteristic densities of AMPA receptors and Ca_V_2.1 channels were determined based on the nearest-neighbor distances (NNDs) between immunogold particles, their clustering properties, and the center-periphery index (CPI), as calculated by Darea following previously described methods.^27^ In this software, Monte-Carlo simulations were used to generate random or fitted particle distributions to evaluate the significance of clustering and center-periphery preferences in subsynaptic localization by comparing empirical data (real) with simulated datasets (sim). The particle clusters were defined by two criteria: the minimum number of particles required to form a cluster and the maximum allowed distance between particles within the same cluster. We specified a minimum of three particles to form a cluster and set the maximum inter-particle distance to the mean NND plus two times its standard deviation.

### Conventions and statistics

All values were reported as the mean ± standard error of the mean (Mean ± SEM) unless otherwise specified. Data sets were tested for normal distribution using the Kolmogorov-Smirnov test. Statistical significance between two groups with unequal variances was tested using an unpaired t-test with Welch’s correction. For comparisons involving more than two groups and normally distributed data, a 2-way ANOVA with Fisher’s exact post hoc test was used followed by Holm-Šidák multiple comparisons correction. If any assumptions for the Kolmogorov-Smirnov test were violated, the non-parametric Kruskal-Wallis test was used, unless otherwise specified. Differences with *P* < 0.05 were considered significant. In the figures, *, **, ***, and **** denote *P* < 0.05, *P* < 0.01, *P* < 0.001, and *P* < 0.0001, respectively. The coefficient of variation (CV) was defined as the ratio of the standard deviation (SD) to the mean (M); CV = SD / M. To analyze the localization preference of proteins within synaptic specializations (central, peripheral or no preference), we compared the observed center-periphery index (CPI_real_) to simulated data (CPI_sim_).^27^ If the CPI_real_ fell within the lowest 2.5% of the CPI_sim_ distribution, it was classified as a central preference. If the CPI_real_ was within the highest 2.5% range of CPI_sim_, then it was classified as a peripheral preference. In all other cases, no preference was assigned. Throughout the paper, data from non-epileptic control individuals and epilepsy patients were shown in black and magenta, respectively. In all figures, data points from females (♀) were displayed using the geometric shape ●, and data points from males (♂) were represented using the geometric shape ▲.

## Results

We quantified the density and distribution of postsynaptic AMPA receptors and presynaptic Ca_V_2.1 channels at excitatory synapses in layers II/III of the human temporal cortex from patients with TLE and non-epileptic surgical comparison cases hereafter referred to as controls. Synapses were classified as dendritic spine synapses onto pyramidal neurons or shaft synapses onto inhibitory interneurons. We first compared postsynaptic AMPA receptor density and distribution between five controls and six patients with TLE, then analyzed presynaptic Ca_V_2.1 channel density and distribution within the AZs of glutamatergic boutons at the same synapse types in the same individuals.

### 1. Identification of pre- and postsynaptic membrane specializations in human replica samples

Excitatory synapses onto dendritic spines and shafts were identified in the freeze-fracture replica preparations of human cortical tissue based on morphological criteria (Fig. 1A-J). In replica membranes, the postsynaptic specialization corresponding to the postsynaptic density (PSD) was reliably recognized by dense aggregations of intramembrane particles (IMPs) on the E-face (extracellular leaflet, Fig. 1D, H, J), consistent with previous observations in rodents.^28^ We identified the presynaptic active zone (AZ) on the P-face (protoplasmic leaflet) based on the elevated IMP density, concave membrane curvature, and clustered synaptic vesicles visible in cross-fractures (Fig. 1E-F, I-J). Synapses on dendritic spines have a steep curvature of the spine head membrane (Fig. 1A, D-F), whereas those on dendritic shafts have relatively flat membrane on larger tube-like profiles (Figure 1B, H-J).

**Figure 1:**
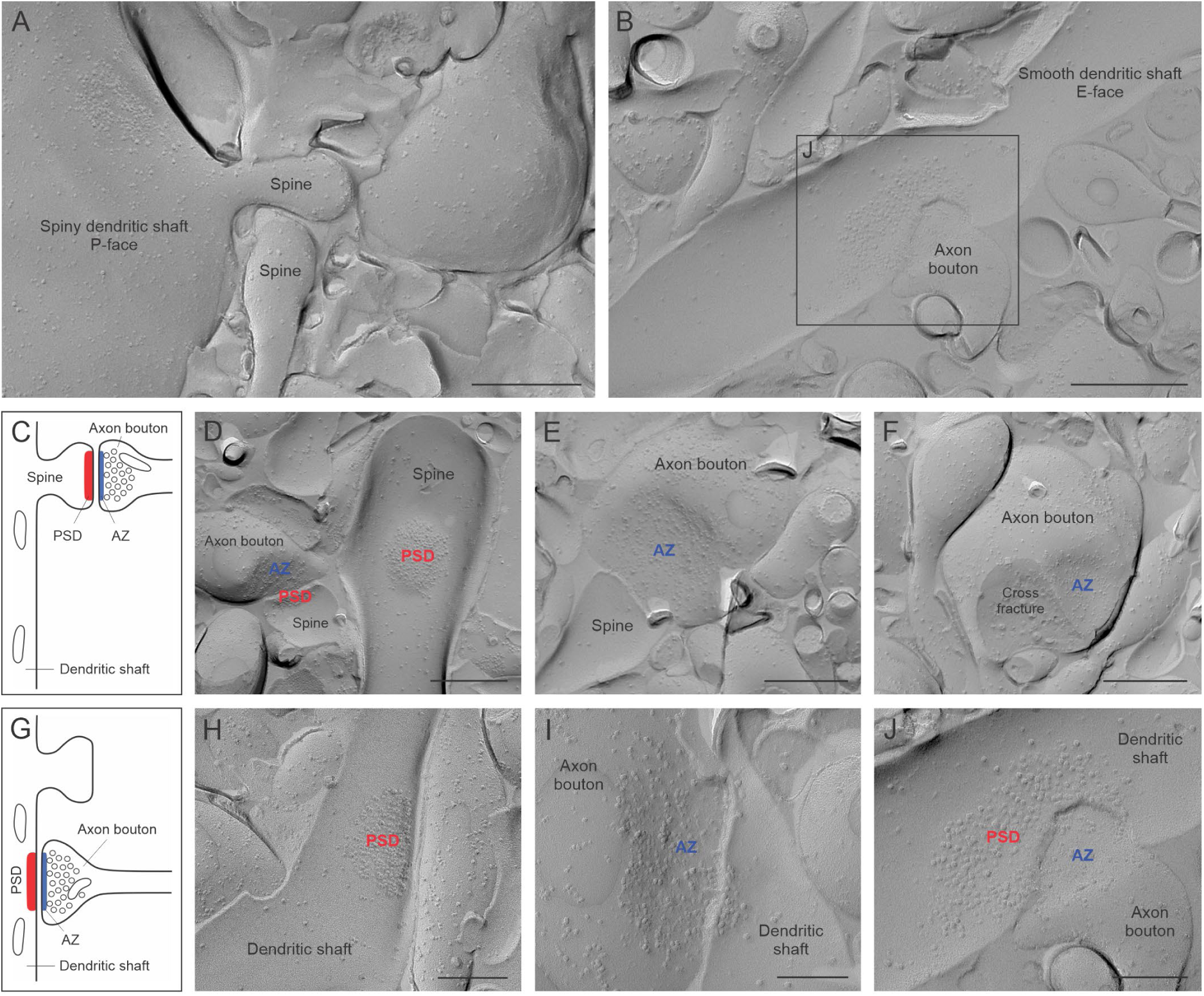
Characterization of dendritic spine and shaft synapses formed by excitatory neurons in replicas of the human temporal cortex. (**A**) Low-magnification image of a spiny pyramidal neuron dendritic shaft showing characteristic intramembrane particle (IMP) density on the P-face and spines emerging from it. Scale bar = 450 nm. (**B**) Low-magnification image of a smooth dendritic shaft from a cortical interneuron with characteristic IMP density on the E-face and an axon bouton forming a synapse with it. Scale bar = 500 nm. (**C**) Schematic illustration of an excitatory dendritic spine synapse with the postsynaptic membrane specialization (PSD) shown in red and the presynaptic active zone (AZ) shown in blue. (**D**) Representative replica image of two dendritic spines displaying dense aggregations of IMPs on the E-face (corresponding to PSDs) and looser aggregations of IMPs on the P-face (corresponding to AZs). Scale bar = 500 nm. (**E**) Replica image of an axon bouton displaying a characteristic IMP density and the concave curvature of the presynaptic P-face membrane, indicating the AZ. Scale bar = 400 nm. (**F**) Replica image of a dendritic spine synapse showing a characteristic P-face specialization marking the AZ, with a cross-fracture plane containing numerous synaptic vesicles. Scale bar = 500 nm. (**G**) Schematic illustration of an excitatory dendritic shaft synapse with the PSD in red and the AZ in blue. (**H**) Replica image of a smooth dendritic shaft showing a dense aggregation of IMPs on the E-face, indicating the PSD. Scale bar = 280 nm. (**I**) Replica image of an axon bouton forming a dendritic shaft synapse, with a characteristic density of IMPs on the P-face and slight membrane curvature that mark the AZ. Scale bar = 200 nm. (**J**) Higher magnification replica image of the dendritic shaft synapse boxed in B, showing the transition from the axonal P-face (dense IMPs) to the dendritic shaft E-face (aggregated IMPs). Scale bar = 200 nm.

### 2. Adapting the deep-learning model *Darea* to human synapse segmentation

To enable high-throughput, minimal-bias analysis of synaptic protein distribution in human replica samples, we adapted the deep learning-assisted image analysis suite Darea, which performs semi-automatic delineation of synaptic membrane specialization.^27^ Because the original model had been trained on rodent replicas, automatic segmentation of human PSDs and AZs required re-training using manual demarcation of these membrane specializations. The demarcation of the PSDs was clear based on structural cues from densely packed IMPs on the E-face (Fig. 2A1-2), while that of AZs was facilitated by simultaneous immunolabeling for the AZ marker proteins RIM1/2, CAST, and neurexin (Fig. 2B1-2, detailed in Supplementary Table 1 Key resources; see also Chen et al.^25^ for reference), along with structural cues. We re-trained the network using manually annotated 200 PSD and 250 AZ profiles in human replica samples. To verify successful retraining, we evaluated performance on different sets of 320 human PSD and 360 human AZ images (taken from non-epileptic control patients) using the original and re-trained versions of Darea (Fig. 2C-D). Re-training substantially improved delineation accuracy for both PSDs and AZs and increased correct detection rates for PSDs from 73% to 95% (Fig. 2C) and AZs from 57% to 93% (Fig. 2D). Minor manual corrections were still occasionally required, but these did not significantly alter the measured areas (Fig. 1C). This optimized Darea workflow was subsequently used to quantitatively analyze synaptic protein organization.

**Figure 2:**
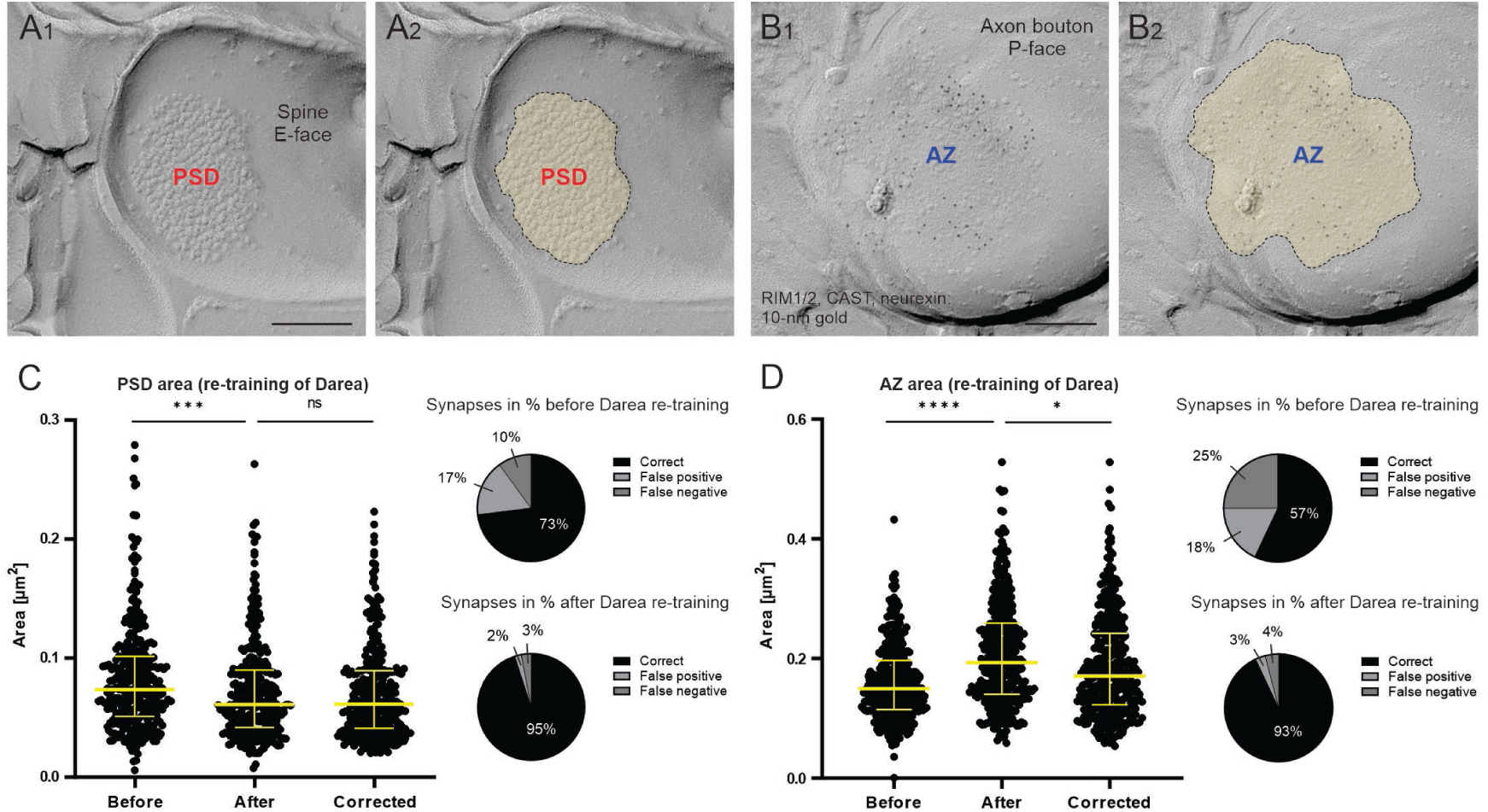
Segmentation of synaptic specializations using Darea. (**A**) Higher magnification replica images of the PSD at a dendritic spine E-face before (A1) and after (A2) manual demarcation, with the PSD area highlighted in yellow. Scale bar = 200 nm. (**B**) Replica images of the presynaptic AZ at the axon bouton P-face immunolabeled for RIM1/2, CAST, and neurexin (10-nm gold each) before (B1) and after (B2) manual demarcation, with the AZ area highlighted in yellow. Scale bar = 150 nm. (**C**) Scatterplot showing the PSD area of individual synapses measured by the deep-learning-assisted replica image analysis suite, Darea (median with interquartile range). Measurements are shown before and after the re-training of the software on human replica samples (200 raw images used). The delineation by Darea was checked by a trained experimenter (Corrected). Statistical differences were analyzed using the Kolmogorov-Smirnov test for nonparametric data. (Before vs. After, *P* = 0.0004; After vs. Corrected, *P* = 0.9847). Proportions of correct, false-positive, and false-negative synapse detections before and after the re-training are shown at right (320 images tested). (**D**) Scatterplot showing AZ areas (median with interquartile range) measured using Darea before and after retraining on human replica samples (250 raw images), with manual validation (Corrected) by a trained experimenter. Statistical differences were analyzed by the Kolmogorov-Smirnov test (Before vs. After, *P* < 0.0001; After vs. Corrected, *P* = 0.0053). Proportions of correctly, falsely positive, and falsely negative detections are shown at right (360 images tested). ****, *P* < 0.0001; ***, *P* < 0.001; **, *P* < 0.01; *, *P* < 0.05; ns, *P* > 0.05, not significant.

### 3. Increased synaptic AMPAR density in excitatory spine synapses in TLE

We next applied this approach to examine the organization of postsynaptic AMPARs at excitatory synapses onto dendritic spines in control and TLE samples. Most of these spine synapses are formed with excitatory pyramidal neurons. Although some inhibitory interneurons also possess dendritic spines, their contribution was not considered here, because spiny non-pyramidal neurons are largely restricted to the middle cortical layers and represent <5% of the total population of cortical neurons.^26,29^ In TLE samples, dendritic spine synapses exhibited a significant increase in AMPAR density compared with non-epileptic controls (Fig. 3A-E). While the overall size of the PSD did not differ between groups (Fig. 3F), the number of AMPAR-associated immunoparticles within the PSD was significantly increased in TLE patients (Fig. 3G). Accordingly, AMPAR density was elevated by approximately 50% in TLE synapses compared with controls (Fig. 3H).

**Figure 3:**
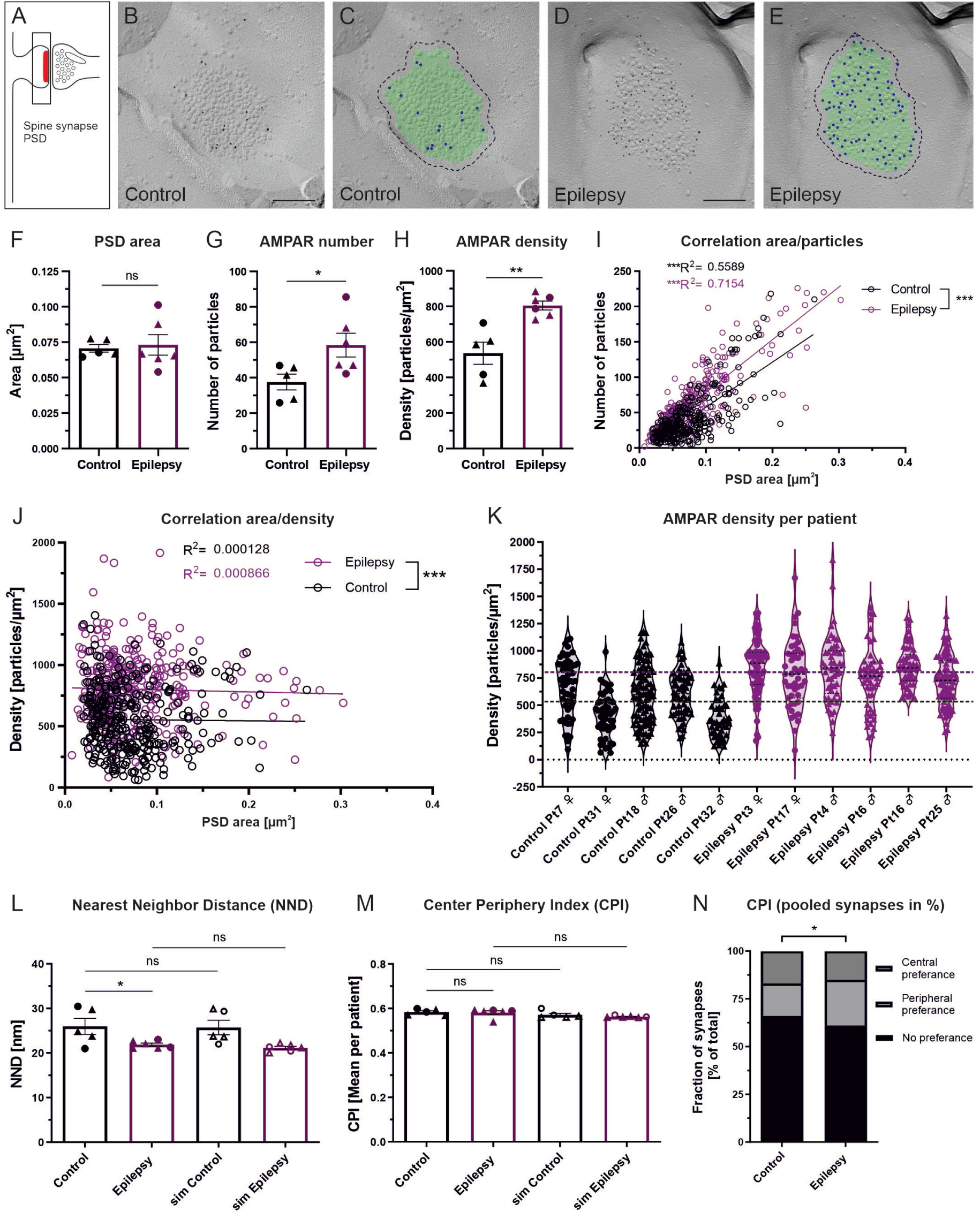
AMPAR localization at dendritic spine synapses in control and TLE patients. (**A**) Schematic illustration of an excitatory dendritic spine synapse with the postsynaptic density (PSD) highlighted in red. (**B**) Representative replica image showing AMPAR immunolabeling (10-nm gold) within the PSD of a dendritic spine synapse from a control patient (Pt18 ♂). Scale bar = 150 nm. (**C**) The same image as in panel B, with the PSD area indicated by a green overlay and immunogold particles marked by blue dots using Darea software. (**D**) Representative replica image showing higher AMPAR immunolabeling within the PSD of a dendritic spine synapse from a TLE patient (Pt3 ♀). Scale bar = 150 nm. (**E**) The same image as in panel D, with the PSD area indicated by a green overlay and immunogold particles marked by blue dots using Darea. (**F**) Bar chart showing PSD areas (mean ± SEM) in control and TLE patients; both complete and incomplete synapses were included in the analysis (Control: 0.073 ± 0.003 µm^2^; Epilepsy: 0.073 ± 0.007 µm^2^; *P* = 0.765; unpaired t-test with Welch’s correction). (**G**) Bar chart showing the mean number of immunogold particles labeling AMPARs in control and TLE patients (Control: 37.6 ± 4.5; Epilepsy: 58.4 ± 6.7; *P* = 0.031; unpaired t-test with Welch’s correction). (**H**) Bar chart showing the mean AMPAR labeling density within PSDs (particles/µm^2^; Control: 536.3 ± 62.80; Epilepsy, 804.9 ± 25.03; *P* = 0.009; unpaired t-test with Welch’s correction). (**I**) Scatterplot illustrating the number of immunoparticles labeling AMPAR relative to the PSD area. All individual synapses are shown for control (n = 324, black) and TLE patients (n = 338, magenta). The number of immunogold particles was positively correlated with PSD area in both control and TLE patients (Control: R^2^ = 0.5589, *P* = 0.0003; TLE: R^2^ = 0.7154; *P* = 0.0003). The regression line was significantly steeper for TLE patients than for controls (*P* = 0.0001; two-way ANOVA with Holm-Šidák correction). (**J**) Scatterplot of AMPAR labeling density (particles/µm^2^) as a function of PSD area. Each data point represents a synapse from control (n = 324, black) or TLE patients (n = 338, magenta). AMPAR labeling density was not correlated with PSD area in either control or TLE patients (Control: R^2^ = 0.000128, *P* = 0.974; TLE: R^2^ = 0.000866; *P* = 0.929), and the slopes of the regression lines did not differ (*P* = 0.975). However, the elevations of the regression lines differed significantly, indicating higher AMPAR immunolabeling density in TLE patients than in controls (*P* = 0.0004; two-way ANOVA with Holm-Šidák correction). (**K**) Violin plot of AMPAR labeling density in PSDs for individual patients. Data sets were normally distributed based on the Kolmogorov-Smirnov test (α = 0.05). The black dashed line indicates the mean density across control patients (M = 535.3 particles/µm^2^; range = 373.5-673.7; CV = 0.26), whereas the magenta dashed line indicates the mean density across TLE patients (M = 800.8 particles/µm^2^; range = 724.8-863.4; CV = 0.076). (**L**) Bar chart of mean nearest-neighbor distances (NNDs) between AMPAR-labeled immunogold particles within PSDs, comparing experimental measurements with Monte Carlo simulations (sim). The experimental NNDs were significantly shorter in the TLE than the control group (Control: 25.9 ± 1.8 nm; Epilepsy: 21.5 ± 0.3 nm; *P* = 0.019; two-way ANOVA with Holm-Šidák correction). Experimental NNDs did not differ significantly from the simulated random distribution in both the control group (*P* = 0.468) and the epilepsy group (*P* = 0.099; two-way ANOVA with Holm-Šidák correction). (**M**) Bar chart of measured Center-Periphery Index (CPI) of AMPARs in spine synapses from control (CPI = 0.577 ± 0.006) and TLE patients (CPI = 0.572 ± 0.008). It did not differ significantly from its corresponding simulated random distribution in the control (Control vs. sim Control: *P* = 0.195) neither in the TLE patients (Epilepsy vs. sim Epilepsy: *P* = 0.0957; two-way ANOVA with Holm-Šidák correction). (**N**) Stacked bar plot of AMPAR CPI categories in pooled synapses from controls and TLE patients. The fraction of synapses showing a central preference was lower in TLE patients than in controls (Control: 17% of total; Epilepsy: 15% of total), whereas the fraction of synapses showing peripheral preference was significantly higher in TLE patients (Control: 17% of total; Epilepsy: 24% of total; *P* = 0.0487; Chi-square test). ●, data points from females (♀); ▴, data points from males (♂); ***, *P* < 0.001; **, *P* < 0.01; *, *P* < 0.05; ns, *P* > 0.05, not significant.

To determine whether this increase was related to synapse size, we analyzed the relationship between PSD area and the number of AMPARs. In both groups, AMPAR number positively correlated with PSD area (Fig. 3I), whereas receptor density was not correlated with PSD area (Fig. 3J), indicating that the increased AMPAR density in TLE patients is independent of synapse size. When we plotted AMPAR density for each individual, the values were normally distributed, with low interindividual variability (Fig. 3K). TLE patients showed significantly higher AMPAR density than the controls, suggesting that AMPAR upregulation occurs consistently in epileptic patients.

We next examined whether AMPARs showed a clustered or random spatial distribution within the PSD. Nearest-neighbor distance analysis revealed significantly shorter distances between AMPAR-associated immunoparticles in TLE synapses than in controls (Fig. 3L), consistent with the increased receptor density observed in TLE. To assess whether receptors formed non-random clusters, we compared the measured nearest-neighbor distances with Monte-Carlo simulated random distributions. No significant differences were detected between experimental and simulated datasets in either group, indicating that AMPARs remained randomly distributed within the PSD.

To quantify the relative location of receptors from the synaptic center toward the PSD border, we next analyzed the center-periphery distribution of AMPARs within the PSD using the center-periphery index (CPI), with values of 0 indicating localization near the PSD center and values of 1 at the PSD border. Average CPI values did not differ significantly between control and TLE patients, nor between experimental and simulated distributions (Fig. 3M). However, pooled analysis of all synapses revealed a modest increase in synapses with peripheral AMPAR localization, consistent with a subtle shift in receptor distribution within the PSD (Fig. 3N). Overall, these analyses demonstrate a robust increase in synaptic AMPAR density in excitatory spine synapses of TLE patients compared with non-epileptic controls.

### 4. Upregulation and reorganization of AMPA receptors in excitatory shaft synapses in TLE patients

To determine whether AMPAR remodeling also occurs at dendritic shaft synapses, we next focused on this distinct population of cortical excitatory synapses. The cellular targets of shaft synapses are more difficult to assign than those of spine synapses, because they are formed predominantly on inhibitory interneurons, but about 10-15% of shaft synapses can also occur on excitatory pyramidal neurons.^26,29,30^ However, we limited our analyses to spine-free dendritic shaft profiles, which originate from inhibitory interneurons. To further assess the potential contribution of pyramidal neuron shaft synapses, we compared AMPAR density between shaft synapses onto spine-free interneurons and those onto pyramidal neurons. We detected no significant difference between these synapse types (Supplementary Fig. 1), suggesting that potential inclusion of pyramidal-neuron shaft synapses is unlikely to substantially affect the overall analysis. Shaft synapses in TLE patients exhibited a pronounced increase in synaptic AMPAR density compared with non-epileptic controls (Fig. 4A-E). Despite the overall similarity in AMPAR upregulation between spine and shaft synapses, shaft synapses exhibited more pronounced changes in receptor density. As in spine synapses, mean PSD area did not differ significantly between groups (Fig. 4F), whereas the number of AMPAR-associated immunoparticles per PSD was significantly increased (Fig. 4G), corresponding to an approximately 70% increase in AMPAR density within the PSD (Fig. 4H). AMPAR-associated particle number positively correlated with PSD area (Fig. 4I), whereas receptor density showed a weak negative correlation with PSD area (Fig. 4J) in both control and TLE groups, indicating relatively higher receptor densities in smaller shaft synapses. Nevertheless, AMPAR densities remained consistently elevated in TLE patients across the full range of synapse sizes. Consistent with the findings in spine synapses, AMPAR density values were normally distributed across individual patients, with low interindividual variability (Fig. 4K). AMPAR densities were significantly higher in TLE patients than in their respective controls. Overall, these findings demonstrate a robust increase in synaptic AMPAR density at excitatory shaft synapses in TLE patients, similar to that observed at spine synapses.

**Figure 4:**
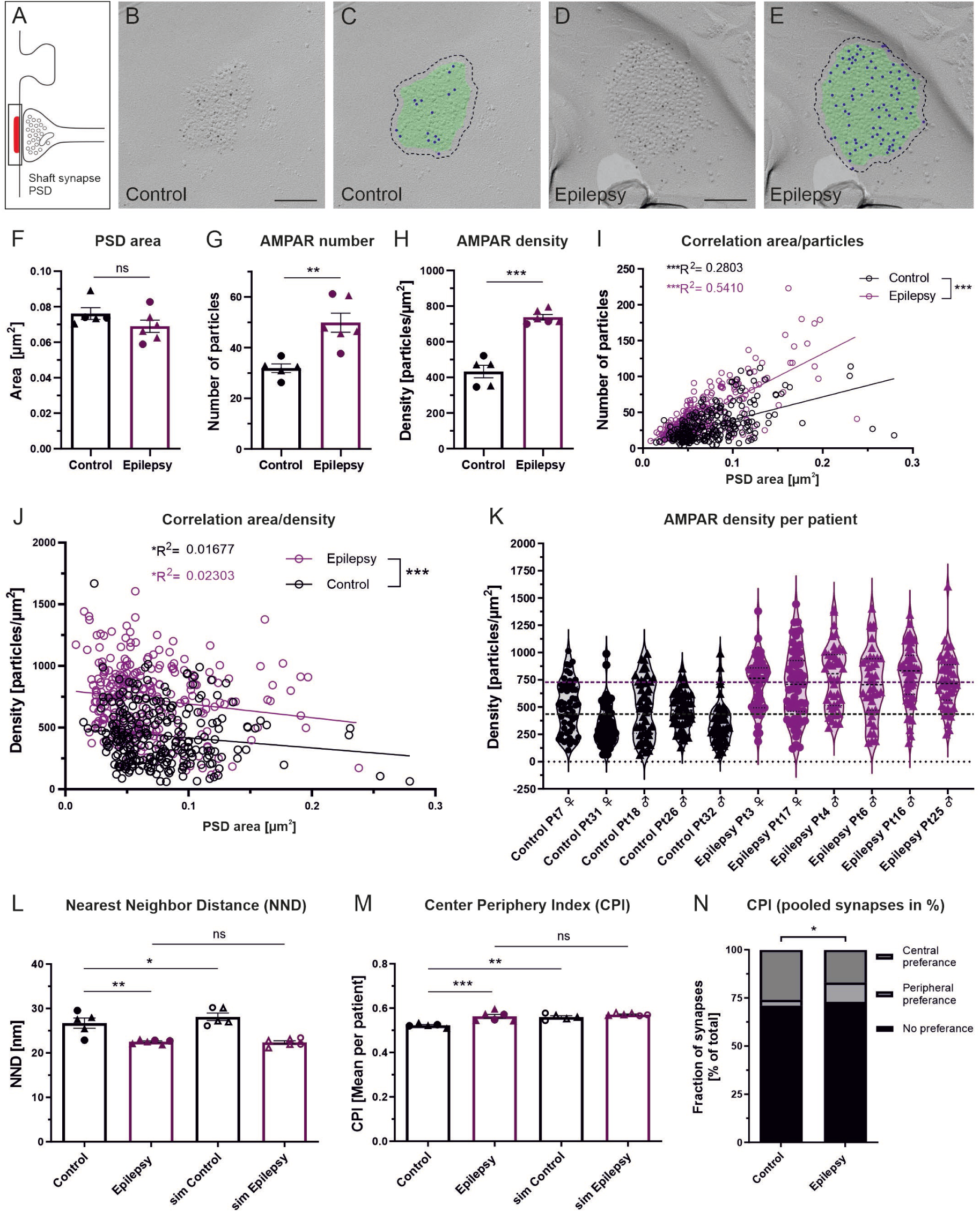
AMPAR localization at dendritic shaft synapses in control and TLE patients. (**A**) Schematic illustration of an excitatory dendritic shaft synapse, with the postsynaptic density (PSD) highlighted in red. (**B**) Representative replica image showing AMPAR immunolabeling (10-nm gold) within the PSD of a dendritic shaft synapse from a control patient (Pt7 ♀). Scale bar = 150 nm. (**C**) The same image as in panel B, with the PSD area indicated by a green overlay and immunogold particles marked by blue dots using the Darea software. (**D**) Representative replica image showing higher AMPAR immunolabeling within the PSD of a dendritic shaft synapse from a TLE patient (Pt25 ♂). Scale bar = 150 nm. (**E**) The same image as in panel D, with the PSD area indicated by a green overlay and immunogold particles marked by blue dots using Darea. (**F**) Bar chart showing PSD areas (mean ± SEM) in control and TLE patients; both complete and incomplete synapses were included in the analysis (Control: 0.076 ± 0.003 µm^2^; Epilepsy: 0.069 ± 0.004 µm^2^; *P* = 0.173; unpaired t-test with Welch’s correction). (**G**) Bar chart showing the mean number of immunogold particles labeling AMPAR in controls and TLE patients (Control: 31.9 ± 1.7; Epilepsy: 49.8 ± 3.8; *P* = 0.004; unpaired t-test with Welch’s correction). (**H**) Bar chart showing the mean AMPAR labeling density within PSDs (particles/µm^2^; Control: 433.0 ± 35.19; Epilepsy, 737.7 ± 15.43; *P* = 0.0003; unpaired t-test with Welch’s correction). (**I**) Scatterplot illustrating the number of immunoparticles labeling AMPARs relative to the PSD area. All individual synapses are shown for control (n = 260, black) and TLE patients (n = 272, magenta). The number of immunogold particles was positively correlated with the PSD area in both control and TLE patients (Control: R^2^ = 0.2803, *P* = 0.0003; TLE: R^2^ = 0.5410; *P* = 0.0003). The regression line was significantly steeper for TLE patients than for controls (*P* = 0.0003; two-way ANOVA with Holm-Šidák correction). (**J**) Scatterplot of AMPAR labeling density (particles/µm^2^) as a function of PSD area. Each data point represents a synapse from control (n = 260, black) or TLE patients (n = 272, magenta). AMPAR labeling density was weakly negatively correlated with PSD area in both control and TLE patients (Control: R^2^ = 0.01677, *P* = 0.0395; TLE: R^2^ = 0.02303; *P* = 0.0118). The slopes of the regression lines did not differ (*P* = 0.609). However, the elevations of the regression lines differed significantly, indicating higher AMPAR immunolabeling density in TLE patients than in controls (*P* = 0.0004; two-way ANOVA with Holm-Šidák correction). (**K**) Violin plot of AMPAR labeling density in PSDs for individual patients. Data sets were normal distributed based on the Kolmogorov-Smirnov test (α = 0.05). The black dashed line indicates the mean density across control patients (M = 433.4 particles/µm^2^; range = 346.4-499.3; CV = 0.18), whereas the magenta dashed line indicates the mean density across TLE patients (M = 737.6 particles/µm^2^; range = 699.7-795.0; CV = 0.051). (**L**) Bar chart of mean nearest-neighbor distances (NNDs) between AMPAR-labeled immunogold particles within PSDs, comparing experimental measurements with Monte Carlo simulations (sim). The experimental NNDs were significantly shorter in the TLE than the control group (Control: 27.4 ± 1.2 nm; Epilepsy: 22.5 ± 0.2 nm; *P* = 0.0012; two-way ANOVA with Holm-Šidák correction). Experimental NNDs differed significantly from the corresponding simulations in the control group (*P* = 0.0219) but not in the epilepsy group (*P* = 0.7424; two-way ANOVA with Holm-Šidák correction). This indicates significant receptor clustering in control patients, which is lost in TLE patients. (**M**) Bar chart of measured Center-Periphery Index (CPI) of AMPARs in shaft synapses from control (CPI = 0.541 ± 0.004) and TLE patients (CPI = 0.567 ± 0.009). While the control group differed significantly from its corresponding simulated random distribution (Control vs. sim Control: *P* = 0.0018), the TLE patient group did not (Epilepsy vs. sim Epilepsy: *P* = 0.264). This indicates significantly different subsynaptic distribution patterns with a greater central preference in controls (*P* = 0.0003; two-way ANOVA with Holm-Šidák multiple comparisons correction. (**N**) Stacked bar plot of AMPAR CPI categories in pooled synapses from controls and TLE patients. The fraction of synapses showing central preference was higher in controls than in TLE patients (Control: 26% of total; Epilepsy: 17% of total), whereas the fraction of synapses showing peripheral preference was significantly higher in TLE patients (Control: 3% of total; Epilepsy: 10% of total; *P* = 0.0138; Chi-square test). ●, data points from females (♀); ▴, data points from males (♂); ***, *P* < 0.001; **, *P* < 0.01; *, *P* < 0.05; ns, *P* > 0.05, not significant.

In contrast to the relatively preserved spatial organization observed in spine synapses, shaft synapses exhibited substantial reorganization of AMPAR distribution in TLE patients. Nearest-neighbor distance analysis revealed significantly shorter distances between AMPAR-associated immunogold particles in TLE synapses than in controls (Fig. 4L), consistent with an increased receptor density in TLE. Comparison with Monte Carlo-simulated random distributions demonstrated significant differences between measured and simulated nearest-neighbor distances in controls, but not in TLE patients (Fig. 4L), indicating a transition from clustered receptor organization in controls (Supplementary Fig. 2A, B) toward a more random distribution in TLE. The cumulative frequency distribution of NNDs indicated shorter NNDs in TLE patients, reflected by a steeper slope rather than a clear leftward shift (Supplementary Fig. 2C). This pattern suggests that gaps between AMPAR clusters are filled in patients with TLE, rather than indicating enhanced receptor packing. Analysis of the CPI further supported this conclusion. Whereas control synapses exhibited significant center-preferred receptor localization relative to simulated random distributions, CPI values in TLE synapses did not differ from those of the simulated distributions (Fig. 4M), suggesting loss of center-preferred organization. Consistently, pooled analysis of all synapses demonstrated significant reduction of center preference in TLE patients compared with controls (Fig. 4N). Together, these findings indicate that excitatory shaft synapses in TLE exhibit not only increased synaptic AMPAR density but also substantial reorganization of receptor spatial organization within the PSD.

### 5. Upregulation and enhanced clustering of Ca_V_2.1 channels in excitatory spine synapses in TLE patients

We next investigated whether presynaptic Ca_V_2.1 channel organization is altered at excitatory synapses onto dendritic spines in patients with TLE. We restricted our analysis to vGluT1-positive excitatory boutons in cortical layers II/III to selectively examine glutamatergic synapses (Fig. 5A-E). The overall area of complete AZs did not differ significantly between control and TLE patients (Fig. 5F). In contrast, the number of Ca_V_2.1-associated immunoparticles per AZ was significantly higher in TLE patients (Fig. 5G), resulting in a corresponding increase in Ca_V_2.1 channel density (Fig. 5H). Ca_V_2.1 particle number showed a moderate positive correlation with AZ size in both groups (Fig. 5I), whereas channel density exhibited a weak negative correlation with AZ area (Fig. 5J), indicating slightly lower densities in larger AZs in both groups. Compared with AMPAR density, Ca_V_2.1 density displayed greater interindividual variability, particularly among patients with TLE (Fig. 5K). To examine this variability in more detail, we stratified the synapse-level data by sex and compared Ca_V_2.1 densities (Fig. 5L). This comparison suggested no apparent sex differences within the control group, but indicated a significant increase in Ca_V_2.1 channel density in male TLE patients, with mean values nearly twice those observed in the controls (Fig. 5L). Overall, these findings suggest increased presynaptic Ca_V_2.1 channel density at excitatory dendritic spine synapses, with this effect appearing more prominent in male patients with TLE.

**Figure 5:**
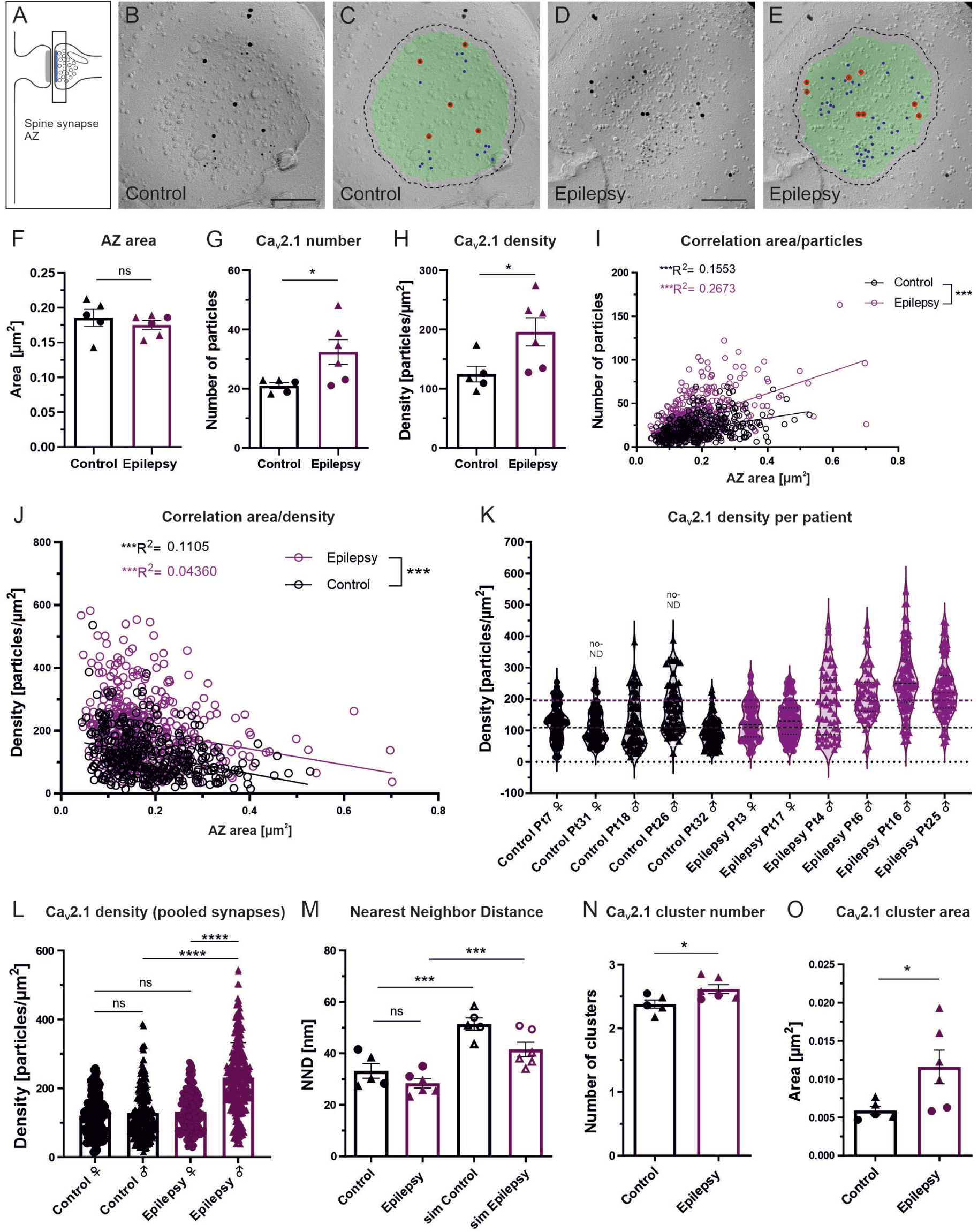
Ca_V_2.1 localization at dendritic spine synapses in control and TLE patients. (**A**) Schematic illustration of an excitatory dendritic spine synapse, with the presynaptic active zone (AZ) highlighted in blue. (**B**) Representative replica image showing immunolabeling for Ca_V_2.1 (10-nm gold) and vGluT1 (15-nm gold) within the AZ of a dendritic spine synapse from a control patient (Pt18 ♂). Scale bar = 150 nm. (**C**) The same image as in panel B, with the AZ area indicated by a green overlay, Ca_V_2.1 immunogold particles marked by blue dots, and vGluT1 immunogold particles marked by red dots using the Darea software. Black dashed line indicates 25 nm outer rim from the demarcation. (**D**) Representative replica image showing higher Ca_V_2.1 immunolabeling within the AZ of a dendritic spine synapse from a TLE patient (Pt16 ♂). Scale bar = 150 nm. (**E**) The same image as in panel D, with the AZ area indicated by a green overlay, Ca_V_2.1 immunogold particles marked by blue dots, and vGluT1 immunogold particles marked by red dots using Darea. Black dashed line indicates 25 nm outer rim from the demarcation. (**F**) Bar chart showing AZ areas (mean ± SEM) in control and TLE patients; only complete synapses were included in the analysis (Control: 0.186 ± 0.012 µm^2^; Epilepsy: 0.175 ± 0.006 µm^2^; *P* = 0.473; unpaired t-test with Welch’s correction). (**G**) Bar chart showing the mean number of immunogold particles labeling Ca_V_2.1 in control and TLE patients (Control: 21.0 ± 1.0; Epilepsy: 32.4 ± 4.2; *P* = 0.041; unpaired t-test with Welch’s correction). (**H**) Bar chart showing the mean Ca_V_2.1 labeling density within AZs (particles/µm^2^; Control: 124.5 ± 13.26; Epilepsy: 196.0 ± 23.96; *P* = 0.032; unpaired t-test with Welch’s correction). (**I**) Scatterplot illustrating the number of immunoparticles labeling Ca_V_2.1 relative to the AZ size. All individual synapses are shown for control (n = 350, black) and TLE patients (n = 424, magenta). The number of immunogold particles was positively correlated with AZ area in both controls and TLE patients (Control: R^2^ = 0.1553, *P* = 0.0003; TLE: R^2^ = 0.2673; *P* = 0.0003). The regression line was significantly steeper for TLE patients than for controls (*P* = 0.0003; two-way ANOVA with Holm-Šidák correction). (**J**) Scatterplot of Ca_V_2.1 labeling density (particles/µm^2^) as a function of AZ area. Each data point represents a synapse from control (n = 350, black) or TLE patients (n = 424, magenta). Ca_V_2.1 labeling density was weakly negatively correlated with AZ area in both controls (Control: R^2^ = 0.1105, *P* = 0.0004) and TLE patients (TLE: R^2^ = 0.0436; *P* = 0.0004). The slopes of the regression lines did not differ (*P* = 0.7266). However, the elevation of the regression lines differed significantly, indicating a higher Ca_V_2.1 immunolabeling density in TLE patients than in controls (*P* = 0.0004; two-way ANOVA with Holm-Šidák correction). (**K**) Violin plot of Ca_V_2.1 labeling density in AZs for individual patients. Data sets were assessed for normal distribution using the Kolmogorov-Smirnov test (α = 0.05). The black dashed line indicates the mean density across control patients (M = 126.8 particles/µm^2^; range = 96.09-168.3; CV = 0.24), whereas the magenta dashed line indicates the mean density across TLE patients (M = 193.3 particles/µm^2^; range = 127.5-262.4; CV = 0.30). (**L**) Combined scatterplot and bar chart showing the mean Ca_V_2.1 labeling density in AZs of pooled synapses from female (♀) and male (♂) control subjects (Control ♀: 120.9 ± 56.39; n = 140 synapses, black; Control ♂: 128.1 ± 74.86; n = 210 synapses, black), and from female (♀) and male (♂) TLE patients (Epilepsy ♀: 131.7 ± 57.74; n = 156 synapses, magenta; Epilepsy ♂: 231 ± 101.4; n = 268 synapses, magenta). Data are shown as mean ± SD. There was a significant difference in Ca_V_2.1 labeling density between female and male Epilepsy patients (*P* < 0.0001), but no significant difference between female and male control patients (*P* > 0.9999). When comparing control and TLE groups, females showed similar density values (*P* > 0.9999), but males exhibited a significantly higher Ca_V_2.1 channel density in the TLE group compared with controls (*P* < 0.0001; Kruskal-Wallis test followed by Dunn’s multiple comparisons test). (**M**) Bar chart of mean nearest-neighbor distances (NNDs) between Ca_V_2.1-labeled immunogold particles within AZs, comparing experimental measurements with Monte Carlo simulations (sim). The experimental NNDs did not differ significantly between the TLE and control groups (Control: 33.2 ± 2.9 nm; Epilepsy: 28.4 ± 1.8 nm; *P* = 0.072; two-way ANOVA with Holm-Šidák correction). In contrast, they differed significantly from the simulated random distribution in both the controls (Control vs. sim Control: *P* = 0.0002) and the TLE patients (Epilepsy vs. sim Epilepsy: *P* = 0.0002; two-way ANOVA with Holm-Šidák correction). This indicates significant channel clustering in both control and TLE patients. (**N**) Bar chart showing the mean Ca_V_2.1 cluster number in spine synapses of control and TLE patients. Patients with TLE disclosed a slight but significant higher number of clusters compared with controls (Control: 2.38 ± 0.06; Epilepsy: 2.62 ± 0.07; *P* = 0.034; unpaired t-test with Welch’s correction). (**O**) Bar chart of Ca_V_2.1 cluster area (mean ± SEM) in spine synapses of control and TLE patients. The cluster area was significantly larger in TLE patients compared with controls (Control: 0.006 ± 0.001 µm^2^; Epilepsy: 0.012 ± 0.002 µm^2^; *P* = 0.048). The larger cluster area was particularly evident in the male patients. ●, data points from females (♀); ▴, data points from males (♂); no-ND, no normal distribution; ****, *P* < 0.0001; ***, *P* < 0.001; **, *P* < 0.01; *, *P* < 0.05; ns, *P* > 0.05, not significant.

We next analyzed the subsynaptic organization of Ca_V_2.1 channels within the AZ. In contrast to the random distribution of AMPARs observed at excitatory spine synapses, nearest-neighbor distance analysis revealed highly significant clustered organization in both control and TLE patients (Fig. 5M). Cluster analysis further revealed a significantly increased number of clusters (Fig. 5N) and cluster size in TLE patients (Fig. 5O). In contrast, center-periphery index (CPI) analysis revealed no significant differences between control and TLE patients or between measured and simulated distribution (Supplementary Fig. 3A), indicating the absence of preferential central or peripheral localization of Ca_V_2.1 channels within the AZ. Together, these findings indicate that excitatory spine synapses in TLE exhibit not only increased Ca_V_2.1 channel density but also enhanced presynaptic Ca_V_2.1 clustering.

### 6. No major changes in Ca_V_2.1 channel density or subsynaptic organization in excitatory shaft synapses in TLE patients

Lastly, we analyzed Ca_V_2.1 channel localization at excitatory synapses onto dendritic shafts in cortical layers II/III (Fig. 6A-E). In contrast to excitatory spine synapses, neither the area of AZs nor the number or density of Ca_V_2.1-associated immunoparticles differed significantly between controls and TLE patients (Fig. 6F-H). Similar to the findings at spine synapses, weak positive correlations were observed between Ca_V_2.1 particle number and AZ size, whereas weak negative correlations were detected between channel density and AZ area in both groups (Fig. 6I-J). Ca_V_2.1 density values showed relatively high interindividual variability in both groups (Fig. 6K), and several patients exhibited non-normal distributions. Overall, these findings indicate that Ca_V_2.1 upregulation in TLE is synapse type-specific and predominantly affects excitatory spine synapses.

**Figure 6:**
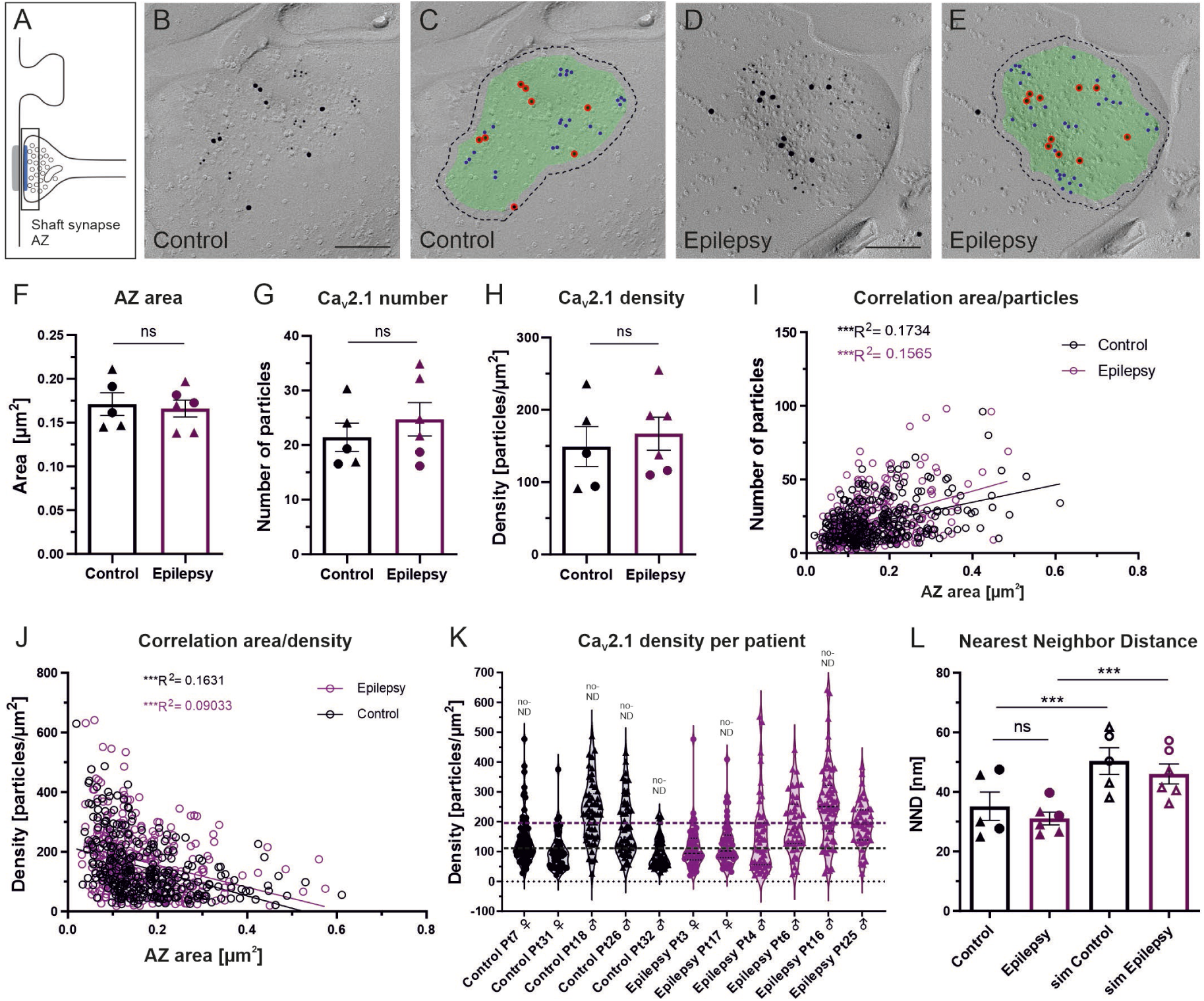
Ca_V_2.1 localization at dendritic shaft synapses in control and TLE patients. (**A**) Schematic illustration of an excitatory dendritic shaft synapse, with the presynaptic active zone (AZ) highlighted in blue. (**B**) Representative replica image showing characteristic Ca_V_2.1 immunolabeling (10-nm gold) within the AZ of a dendritic shaft synapse from a control patient (Pt26 ♂). (**C**) The same image as in panel B, with the AZ area indicated by a green overlay, Ca_V_2.1 immunogold marked by blue dots, and vGluT1 immunogold marked by red dots using the Darea software. Black dashed line indicates 25 nm outer rim from the demarcation. (**D**) Representative replica image displaying similar Ca_V_2.1 immunolabeling within the AZ of a dendritic shaft synapse from a TLE patient (Pt17 ♀). (**E**) The same image as in panel D, with the AZ area indicated by a green overlay, Ca_V_2.1 immunogold marked by blue dots, and vGluT1 immunogold marked by red dots using Darea. Black dashed line indicates 25 nm outer rim from the demarcation. (**F**) Bar chart (mean ± SEM) showing AZ areas in control and TLE patients; only complete synapses were included in the analysis (Control: 0.171 ± 0.013 µm^2^; Epilepsy: 0.166 ± 0.009 µm^2^; *P* = 0.769; unpaired t-test with Welch’s correction). (**G**) Bar chart showing the mean number of immunogold particles labeling Ca_V_2.1 in control and TLE patients (Control: 21.4 ± 2.6; Epilepsy: 24.7 ± 3.0; *P* = 0.431; unpaired t-test with Welch’s correction). (**H**) Bar chart showing the mean Ca_V_2.1 labeling density within AZs (particles/µm^2^; Control: 149.2 ± 27.63; Epilepsy: 167.1 ± 22.86; *P* = 0.631; unpaired t-test with Welch’s correction). (**I**) Scatterplot illustrating the number of immunoparticles labeling Ca_V_2.1 relative to the AZ size. All individual synapses are shown for control (n = 266, black) and TLE patients (n = 299, magenta). The number of immunogold particles was positively correlated with AZ area in both controls (Control: R^2^ = 0.1734, *P* = 0.0004) and TLE patients (Epilepsy: R^2^ = 0.1565; *P* = 0.0004), and the slopes of the regression lines did not differ significantly (*P* = 0.1242; two-way ANOVA with Holm-Šidák correction). (**J**) Scatterplot of Ca_V_2.1 labeling density (particles/µm^2^) as a function of the AZ area. Each data point represents a synapse from control (n = 266, black) or TLE patients (n = 299, magenta). Ca_V_2.1 labeling density was weakly negatively correlated with AZ area in both control (Control: R^2^ = 0.1631, *P* = 0.0004) and TLE patients (Epilepsy: R^2^ = 0.0903; *P* = 0.0004). No significant difference was observed in the slope of the regression lines (*P* = 0.6348) neither in the elevation of regression lines (*P* = 0.0516; two-way ANOVA with Holm-Šidák correction) indicating similar Ca_V_2.1 labeling densities in control and TLE patients. (**K**) Violin plot of Ca_V_2.1 labeling densities in AZs for individual patients. Data sets were assessed for normal distribution using the Kolmogorov-Smirnov test (α = 0.05). The black dashed line indicates the mean density across control patients (M = 152.5 particles/µm^2^, range = 95.15-235.9; CV = 0.41), whereas the magenta dashed line indicates the mean density across TLE patients (M = 171.3 particles/µm^2^, range = 113.2-255.0; CV = 0.34). (**L**) Bar chart of mean nearest-neighbor distances (NNDs) between Ca_V_2.1-labeled immunogold particles within AZs, comparing experimental measurements with Monte Carlo simulations (sim). The experimental NNDs did not differ significantly between the TLE and control groups (Control: 35.2 ± 4.7 nm; Epilepsy: 31.1 ± 2.1 nm; *P* = 0.439; two-way ANOVA with Holm-Šidák correction). In contrast, they differed significantly from the simulated random distribution in both the controls (Control vs. sim Control: *P* = 0.0002) and the TLE patients (Epilepsy vs. sim Epilepsy: *P* = 0.0002; two-way ANOVA with Holm-Šidák correction). This indicates significant channel clustering in both control and TLE patients. ●, data points from females (♀); ▴, data points from males (♂); no-ND, no normal distribution. ***, *P* < 0.001; **, *P* < 0.01; *, *P* < 0.05; ns, *P* > 0.05, not significant.

We next examined the subsynaptic organization of Ca_V_2.1 channels within shaft synapse AZs. Nearest-neighbor distance analysis revealed significant clustering of Ca_V_2.1 channels in both control and TLE patients (Fig. 6L). However, no significant differences in the number of Ca_V_2.1 clusters (Supplementary Fig. 4A) or cluster size were detected between control and TLE groups (Supplementary Fig. 4B), indicating that clustered Ca_V_2.1 organization is a general feature of shaft synapse AZs independent of disease state. Likewise, center-periphery index analysis revealed no significant difference between control and TLE groups or between measured and simulated distributions (Supplementary Fig. 3B), indicating the absence of preferential central or peripheral Ca_V_2.1 localization within AZs of shaft synapses. Together, these findings suggest that, unlike excitatory spine synapses, excitatory shaft synapses do not undergo major disease-associated reorganization of presynaptic Ca_V_2.1 channels in TLE.

## Discussion

Our study combines high-resolution freeze-fracture replica immunolabeling with quantitative nanoscale analysis to investigate the organization of postsynaptic AMPARs and presynaptic Ca_V_2.1 channels at excitatory synapses in human temporal cortex under epileptic and non-epileptic conditions. Our data reveal synapse-type-specific nanoscale reorganization of glutamatergic synapses in TLE. Postsynaptic AMPAR density was elevated at both spine and shaft synapses, whereas presynaptic Ca_V_2.1 reorganization was largely restricted to spine synapses, where channels showed enhanced clustering within active zones. By contrast, shaft synapses exhibited no detectable changes in Ca_V_2.1 organization. Together, these findings identify distinct forms of pre- and postsynaptic remodeling that may contribute to cortical hyperexcitability and altered excitatory transmission.

### Postsynaptic modulation of AMPAR density and distribution

The most consistent alteration observed in TLE synapses was the marked increase in postsynaptic AMPAR density. Both excitatory dendritic spine and shaft synapses exhibited substantially elevated AMPAR levels despite normal ultrastructure of synaptic specializations (Fig. 1) and unchanged PSD size (Fig. 3 and 4). This is consistent with ultrastructural reports of largely preserved dendritic arborization and spine morphology, even in patients with a long-lasting history of epilepsy and high seizure frequency, and does not support the view that epilepsy necessarily produces a dendritic pathology.^31,32^ Rather, our results suggest that TLE-associated changes primarily involve the molecular composition and nanoscale organization of synaptic specializations.

Notably, the increase in AMPAR density was highly reproducible across patients and showed remarkably low interindividual variability. This increase in receptor density could enhance excitatory synaptic transmission and may thereby contribute to the persistent hyperexcitability characteristic of epileptic cortical circuits.^8,10,15^ Interestingly, AMPAR spatial organization was differentially affected. Spine synapses largely preserved their receptor distribution patterns, whereas shaft synapses displayed pronounced reorganization from clustered to more random arrangements, together with reduced center preference within the PSD. Our data indicate that the smaller NNDs accompanying increased receptor density are more consistent with the filling of gaps between receptor aggregates that with tighter packing with discrete clusters. These observations suggest that postsynaptic remodeling in TLE is not uniform across excitatory synapse populations but instead involves synapse-type-specific nanoscale adaptations that may differentially influence cortical circuit function. In addition, our findings do not support a nano-column hypothesis,^33,34^ but instead are consistent with the comparatively random AMPAR distributions reported at rodent central synapses using similar methodological approaches.^35,36^

### Synapse-type-specific CaV2.1 reorganization

Synapse-type-dependent alterations in synaptic proteins in TLE were most pronounced for the presynaptic voltage-gated calcium channel Ca_V_2.1. The significant increases in Ca_V_2.1 channel density and clustering were largely confined to excitatory spine synapses, whereas shaft synapses exhibited comparatively modest changes. These findings suggest that distinct excitatory synapse populations undergo partially divergent forms of pathological plasticity during chronic epileptic activity. Given that spine and shaft synapses are associated with different postsynaptic targets and circuit functions, this differential nanoscale remodeling may contribute to selective changes in cortical information processing and the excitation-inhibition (E/I) balance in TLE.^37^

Most excitatory spine synapses in the cerebral cortex target pyramidal cells, the principal excitatory projection neurons of the cortex, which play central roles in cortical circuit function, synaptic integration, and information flow.^26,32^ Pyramidal cells are abundant across cortical layers II-VI and account for approximately 70-80% of cortical neurons. In contrast, excitatory shaft synapses often target inhibitory interneurons, which are typically short-axon cells with smooth or sparsely spiny dendrites and account for approximately 15-30% of cortical neurons.^26^ However, postsynaptic cell-type assignment is more challenging for shaft synapses, and a considerable proportion of these synapses also contact excitatory pyramidal neurons.^30^

Despite this heterogeneity in postsynaptic targets, presynaptic Ca_V_2.1 remodeling may selectively alter excitatory drive onto excitatory but not inhibitory neurons in TLE, thereby contributing to a disrupted cortical E/I balance. This interpretation is consistent with the prevailing view that epilepsy involves recurrent seizures arising from abnormal synchronization and disrupted excitatory-inhibitory balance within neuronal networks.^8,37^ Because modulation of excitatory and inhibitory neurotransmission is a central principle of antiseizure pharmacotherapy,^38^ Ca_V_2.1 channels may represent a potential therapeutic target. However, their therapeutic relevance requires further investigation, given their broad roles in neurotransmitter release and normal circuit function. The cellular and circuit mechanisms underlying seizure generation at epileptic foci remain incompletely understood, and even subtle structural and functional alterations may contribute to epileptogenesis.

### Evidence for synaptic plasticity in TLE

Experimental studies indicate that Ca_V_2.1 channels play important roles in cortical neurons and corticothalamic circuits, where they contribute to multiple developmental processes including activity-dependent structural plasticity. Mutations in the *CACNA1A* gene encoding the Ca_V_2.1 α1 subunit, and loss-of-function (LoF) variants, are associated with a range of neurological disorders, including epilepsy. The long-term mechanisms through which biallelic *CACNA1A* LoF variants contribute to epilepsy, including absence and motor seizures, remain incompletely understood.^39^ By contrast, short-term synaptic plasticity is better investigated. Ca_V_2.1-dependent synaptic release contributes to short-term facilitation, whereas loss of Ca_V_2.1 channel function can lead to synaptic depression or reduced facilitation.^17^ Although Ca_V_2.1 dysfunction may contribute to epileptogenesis through altered presynaptic calcium signaling and circuit development, its long-term effects on synaptic plasticity in TLE remain incompletely defined. By contrast, evidence for persistent plasticity-related changes at the postsynaptic level, particularly involving AMPARs, is more direct.

Evidence from experimental models and studies of human TLE suggests that the disease is associated with persistent, potentially maladaptive alterations in AMPAR expression and AMPAR-dependent synaptic plasticity.^40^ In rodents, pilocarpine-induced status epilepticus has been associated with increased cortical expression of AMPAR subunits, consistent with sustained seizure-associated remodeling of excitatory synapses.^8,41^ Complementing these observations, altered AMPAR abundance has been reported in surgical tissue from patients with TLE at the mRNA and protein levels, as well as rodent models of chronic TLE.^42,43^ Importantly, *in vivo* evidence of altered AMPAR availability in TLE has been obtained using PET with the carbon-11-labeled AMPAR tracer ^11C^K-2. In individuals with TLE, ^11C^K-2 uptake was increased and correlated with enhanced regional AMPAR protein expression in corresponding surgical specimens.^44^

Together, these data are consistent with region-specific changes in AMPAR expression and localization. Such changes, together with the increased Ca_V_2.1 in spine synapses, may enhance AMPAR-mediated excitatory transmission and thereby contribute to network hyperexcitability, epileptogenesis, or seizure persistence.^8^ However, it is more likely that these changes reflect impaired homeostatic plasticity rather than primary causal link, considering the divergent etiology, clinical history, and medication in TLE patients. The available studies also do not establish a direct causal relationship. Consistent with a functional contribution of excessive AMPAR signaling, preclinical studies indicate that pharmacological AMPAR antagonism or interventions targeting synaptic AMPARs can reduce seizure activity in experimental epilepsy models.^45^ However, AMPAR regulation differs across brain regions, disease stages, cell types, and receptor subunit compositions, and further work is required to distinguish causally epileptogenic alterations from compensatory adaptations in TLE.

### A trend toward sex-dependent effects

We observed a potential sex-dependent difference in presynaptic Ca_V_2.1 remodeling at dendritic spine synapses (Fig. 5). Increased Ca_V_2.1 density in TLE samples was detected primarily in male patients, whereas samples from female patients showed no comparable change. Although this observation requires cautious interpretation, it suggests that presynaptic calcium-channel regulation at pyramidal cell synapses may be influenced by sex-dependent molecular or hormonal factors.^46^ Conversely, altered synaptic Ca_V_2.1 expression could contribute to seizure susceptibility, disease progression, and treatment response. Sex-related differences in epilepsy phenotypes, seizure susceptibility, and clinical presentation have been reported with higher rates of excitability episodes and seizures in men than in women. However, these relationships are complex and may reflect interacting biological, clinical, and social factors.^47^ Experimental evidence for sex-specific modulation of excitatory neurotransmission supports the biological plausibility of this hypothesis.^48,49^

Sex-related differences have also been reported in responses to antiseizure medications (ASMs), although their mechanisms and clinical relevance remain incompletely understood.^47^ Differences in hormone signaling, neural circuitry, and synaptic molecular composition may contribute to variable seizure vulnerability and treatment responses. A recent review found that most studies did not identify significant sex differences in ASM efficacy. However, among studies reporting an association, several described better outcomes in males, including higher rates of seizure reduction or seizure freedom.^50^

The basis of these differences remains unclear, but sex-dependent variation in the molecular composition of synaptic specializations represents a plausible mechanism. Larger, well-characterized patient cohorts are needed to determine whether the Ca_V_2.1 differences observed in our study represent robust biological sex differences in TLE. Future studies should also examine whether sex-dependent synaptic molecular composition contributes to epileptogenesis or ASM responsiveness.

### Technical considerations

The SDS-FRL method enables high-resolution mapping of transmembrane proteins and offers several advantages over other EM immunolabeling techniques, including high sensitivity, reproducibility, and quantitative analysis.^22,51,52^ Furthermore, to our knowledge, this study represents the first application of the SDS-FRL method to investigate synaptic proteins in human tissue. Nevertheless, the approach remains primarily structural and does not directly assess synaptic physiology, release probability, or channel function. Future studies combining ultrastructural analyses with electrophysiological and functional imaging approaches will therefore be important for establishing direct relationships between nanoscale molecular organization and functional alterations at excitatory synapses in epileptic circuits.^53,54^ In addition, larger cohorts will be necessary to account for potentially confounding clinical variables, including disease duration, medication history, and seizure burden.

## Conclusions

In summary, our study shows that temporal lobe epilepsy is associated with distinct forms of molecular nanoscale remodeling at excitatory synapses in the human temporal cortex. Increased postsynaptic AMPAR density at both spine and shaft synapses suggests enhanced excitatory signaling capacity, whereas selective reorganization of presynaptic Ca_V_2.1 localization at spine synapses indicates that distinct synapse populations may engage in different forms of pathological plasticity. These findings provide ultrastructural evidence that pathological plasticity in epilepsy involves differential and potentially target-cell-specific alterations in pre- and postsynaptic nanoarchitecture. More broadly, the present work highlights nanoscale synaptic organization as a potential contributor to excitatory circuit dysfunction in temporal lobe epilepsy and a promising focus for future mechanistic studies and therapeutic strategies.

## Data availability

Further information and requests for resources should be directed to the Lead Contact, Walter A. Kaufmann, who will respond to reasonable requests. Original data are stored in the scientific repositories of ISTA and are available upon request.

## Supporting information

Supplemental Table 1 Key resources

## Acknowledgements

The authors thank Andreas Gruber (Johannes Kepler University Linz and Kepler University Hospital Linz, Department of Neurosurgery) for overseeing the surgical procedures; Alexandra Lang (Medical University of Vienna, Department of Neurosurgery) for substantial support with sample recruitment, procedural coordination, and facilitating access to relevant patient information; and David Vandael (Institute of Anatomy and Cell Biology, Johannes Kepler University Linz) for his assistance with sample recruitment. This research was supported by the Scientific Service Units (SSUs) of ISTA through resources provided by the Electron Microscopy Facility (EMF).

## Supplementary Material

The supplementary material comprises the following files:

- Supplementary Table 1 Key resources
- Supplementary Figures 1-4

