## Supplemental Table 1 Key resources for "Nanoscale remodeling of synaptic glutamate receptors and calcium channels in temporal lobe epilepsy"

### Supplementary Table 1: Key resources

| Reagent or Resource | Source | Identifier |
|---|---|---|
| **Antibodies** | | |
| Guinea pig Polyclonal anti-pan-AMPAR | Frontier Institute co.,ltd. | RRID: AB_2571610 |
| Guinea pig Polyclonal anti-VGCC (Cav2.1) α-1A | Synaptic Systems | Cat # 152 205 |
| Rabbit Polyclonal anti-VGLUT 1 | Synaptic Systems | Cat # 135 303 |
| Rabbit Polyclonal anti-RIM 1/2 | Synaptic Systems | Cat # 140 203 |
| Rabbit Polyclonal anti-CAST 1 | Provided by Akari Hagiwara and Toshihisa Ohtsuka, Yamanashi University Medical School, Japan | NP_740768 |
| Rabbit Polyclonal anti-pan-Neurexin | Provided by Masahiko Watanabe; Hokkaido University School of Medicine, Japan | N/A |
| Goat anti-guinea pig IgG:5 nm immunogold conjugate | British Biocell International (BBI), Cardiff | Cat # EM.GAG5 |
| Goat anti-guinea pig IgG:10 nm immunogold conjugate | British Biocell International (BBI), Cardiff | Cat # EM.GAG10 |
| Goat anti-rabbit IgG:5 nm immunogold conjugate | British Biocell International (BBI), Cardiff | Cat # EM.GAR5 |
| Goat anti-rabbit IgG:10 nm immunogold conjugate | British Biocell International (BBI), Cardiff | Cat # EM.GAR10 |
| Goat anti-rabbit IgG:15 nm immunogold conjugate | British Biocell International (BBI), Cardiff | Cat # EM.GAR15 |
| **Chemicals and recombinant proteins** | | |
| Bovine serum albumin (BSA) | Sigma-Aldrich | Cat # A9647 |
| Chloroform | Sigma-Aldrich | Cat # 372978 |
| Durcupan ACM Resin Single component A | Sigma-Aldrich | Cat # 44611 |
| Durcupan ACM Resin Single component B | Sigma-Aldrich | Cat # 44612 |
| Durcupan ACM Resin Single component C | Sigma-Aldrich | Cat # 44613 |
| Durcupan ACM Resin Single component D | Sigma-Aldrich | Cat # 44614 |
| Glutaraldehyde | Electron Microscopy Sciences | Cat # 16220 |
| Glycerol | Sigma-Aldrich | Cat # G5516 |
| Glycine | VWR | Cat # 24403.298 |
| KCl | VWR | Cat # 26764.232 |
| Lead (II) nitrate | Sigma-Aldrich | Cat # 228621 |
| NaCl | Sigma-Aldrich | Cat # S7653 |
| $Na_2HPO_4$ | VWR (Merck) | Cat # 1.06580 |
| $NaH_2PO_4$ | VWR (Merck) | Cat # 1.06346 |
| NaOH | Sigma-Aldrich | Cat # S5881 |
| Paraformaldehyde | TAAB | Cat # FO 17/1 |
| Picric acid solution (saturated) | Sigma-Aldrich | Cat # P6744 |
| Polyethylene glycol 2000 | VWR (Merck) | Cat # 8.21037 |
| Polyvinyl Formal Resin 15/95 | Electron Microscopy Sciences | Cat # 15800 |
| Sodium azide | Sigma-Aldrich | Cat # S2002 |
| Sodium dodecyl sulfate (SDS) | Sigma-Aldrich | Cat # L3771 |
| Sucrose | Sigma-Aldrich | Cat # S0389 |
| Tannic acid | Sigma-Aldrich | Cat # 403040 |
| Tris(hydroxymethyl)amino methane | VWR (Merck) | Cat # 1.08387 |
| Tri-sodium citrate | Sigma-Aldrich | Cat # S1804 |
| Uranyl acetate | Serva | Cat # 77870.02 |
| **Software and algorithms** | | |
| Adobe Photoshop | Adobe Inc. | https://www.adobe.com/products/photoshop.html |
| CorelDRAW 2019 | CorelDRAW | https://www.coreldraw.com/en |
| Fiji (ImageJ version 1.53c) | NIH, Open source | https://fiji.sc/ |
| MATLAB 2020 | MathWorks | https://www.mathworks.com/ |
| Radius Software | EMSIS | https://www.emsis.eu/products/software/radius/ |
| **Others** | | |
| Aluminum platelets (3 mm OD, 0.1 / 0.2 mm indentation) | Engineering Office M. Wohlwend GmbH, Switzerland | Cat # 241 |
| Aluminum platelets (3 mm OD, 0.3 mm indentation) | Engineering Office M. Wohlwend GmbH, Switzerland | Cat # 242 |
| Copper carriers with flat surface (4.6/3 x 0.6 mm) | Engineering Office M. Wohlwend GmbH, Switzerland | Cat # 528 |
| 100-line/inch Parallel bar copper EM grids | Science Services | Cat # G100P-Cu |
