## Supplemental Figure 1 for "Nanoscale remodeling of synaptic glutamate receptors and calcium channels in temporal lobe epilepsy"

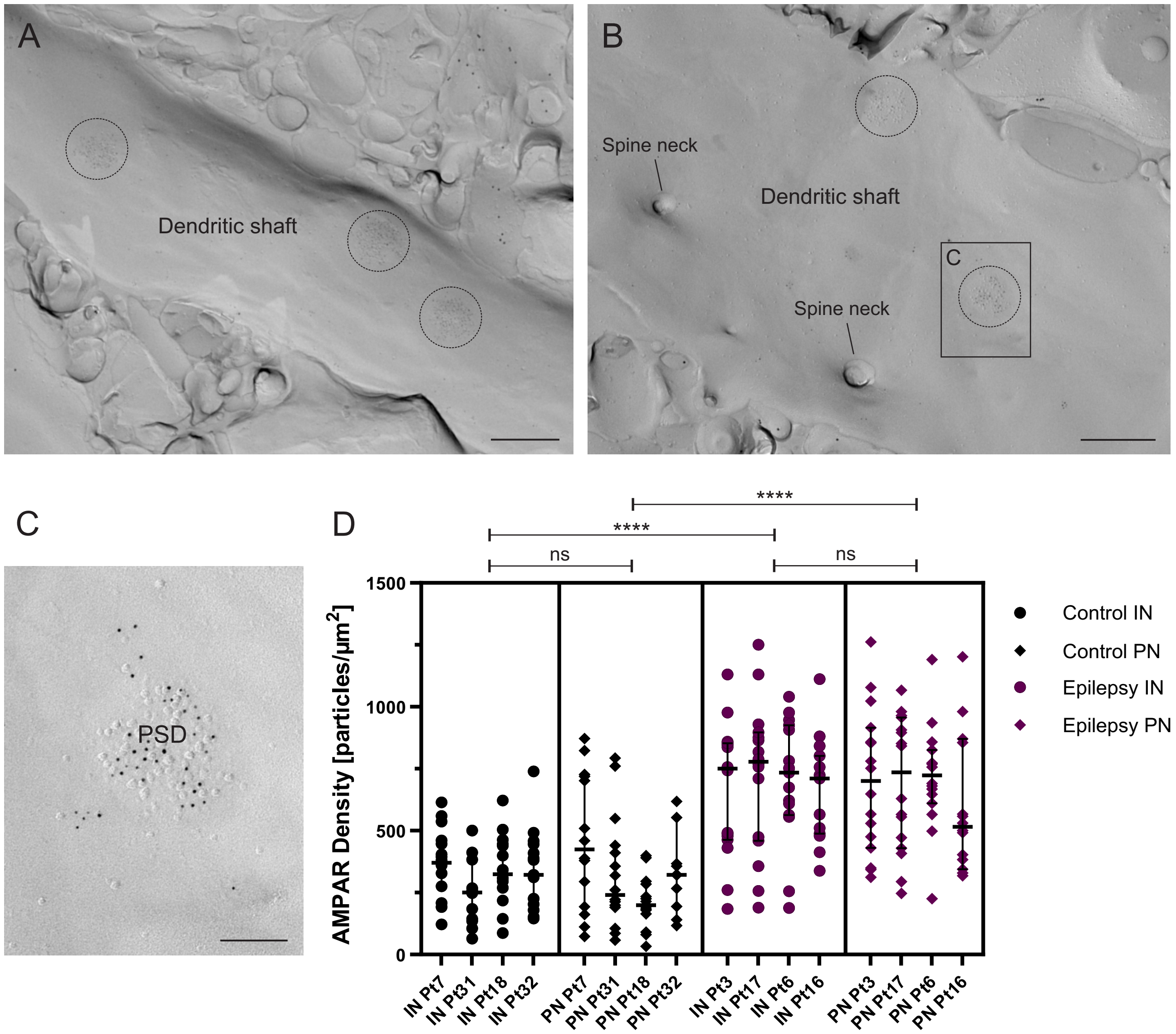


**Supplementary Figure 1: AMPAR immunolabeling density at dendritic shaft synapses of interneurons (INs) and pyramidal neurons (PNs).** (**A**) Replica image of a smooth dendritic shaft from a cortical IN. Postsynaptic densities (PSDs) of excitatory shaft synapses are outlined by dashed circles. AMPAR labeling, 5-nm gold; vGluT1 labeling, 15-nm gold; scale bar = 500 nm. (**B**) Replica image of a spiny dendritic shaft from a cortical PN. PSDs of excitatory shaft synapses are outlined by dashed circles. AMPAR labeling, 5-nm gold; vGluT1 labeling, 15-nm gold; scale bar = 500 nm. (**C**) Higher magnification image of the PN dendritic shaft synapse boxed in B, showing characteristic AMPAR labeling (5-nm gold) within the PSD; scale bar = 300 nm. (**D**) Nested one-way ANOVA (experimental data plus median with 95% CI) showing the AMPAR immunogold labeling density within PSDs at IN shaft synapses in controls (Control IN: median = 321.5; n = 60 synapses; patients included: Pt7 ♀, Pt31 ♀, Pt18 ♂, Pt32 ♂) and PN shaft synapses in controls (Control PN: median = 267.3, n = 55 synapses; patients included: Pt7 ♀, Pt31 ♀, Pt18 ♂, Pt32 ♂). The labeling density did not differ significantly comparing both groups ($P > 0.999$). Likewise, the AMPAR labeling density was similar in IN shaft synapses of TLE patients (Epilepsy IN: median = 738.8, n = 62 synapses; patients included: Pt3 ♀, Pt17 ♀, Pt6 ♂, Pt16 ♂) and PN shaft synapses of TLE patients (Epilepsy PN: median = 658.0, n = 62 synapses; patients included: Pt3 ♀, Pt17 ♀, Pt6 ♂, Pt16 ♂). Again, the labeling density did not differ significantly comparing both groups ($P = 0.997$). However, the overall AMPAR labeling density was significantly higher in TLE patients than in controls for both IN shaft synapses and PN shaft synapses ($P < 0.0001$ for each comparison). Statistics: Nested one-way ANOVA. ****, $P < 0.0001$; ns, $P > 0.05$, not significant.
