## Supplemental Figure 2 for "Nanoscale remodeling of synaptic glutamate receptors and calcium channels in temporal lobe epilepsy"

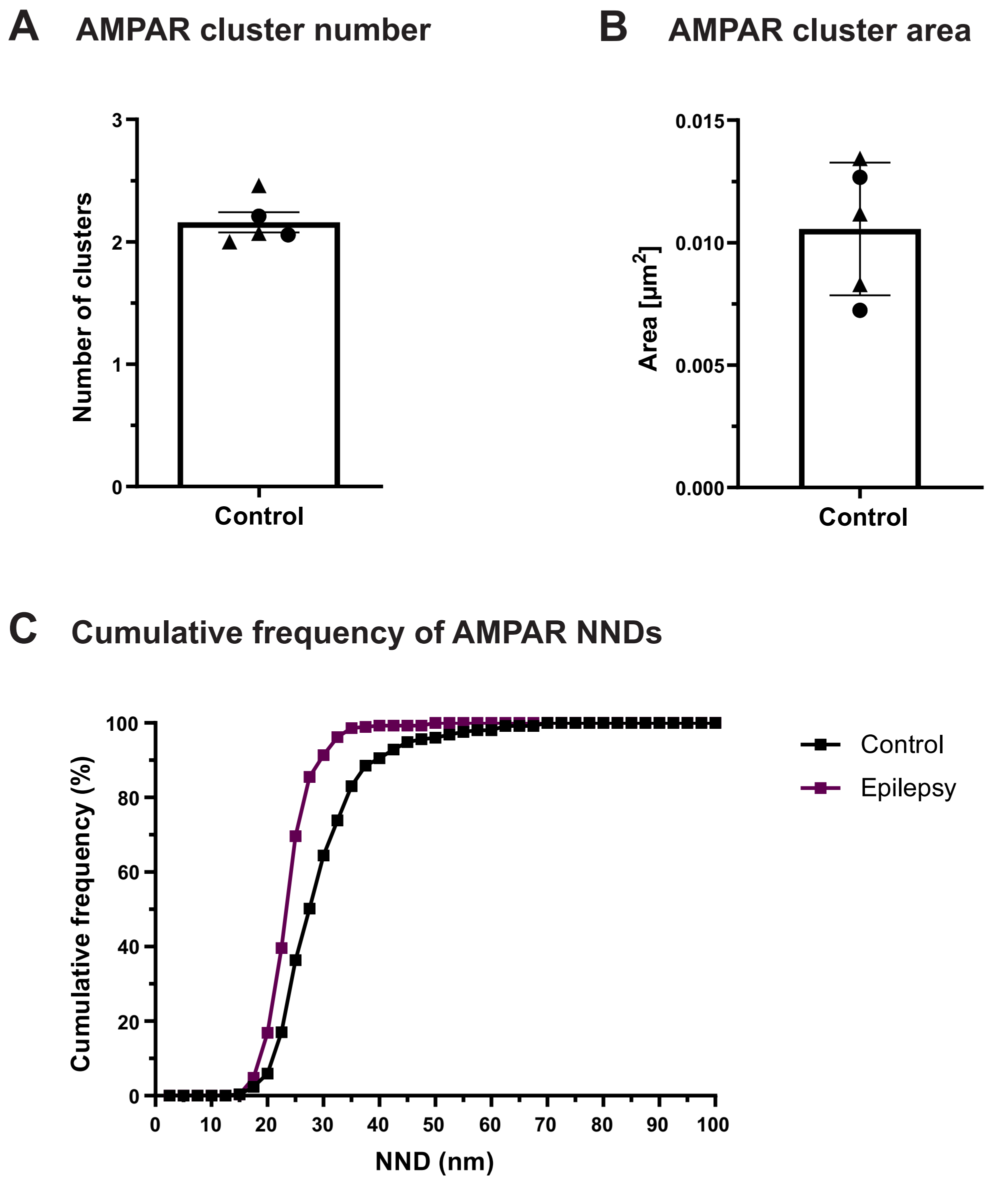


**Supplementary Figure 2: Spatial organization of AMPA receptors within PSDs of dendritic shaft synapses.** (**A**) Bar chart (mean ± SEM) showing the number of AMPAR clusters within PSDs in control samples (Control: n of clusters per PSD = 2.16 ± 0.08). (**B**) Bar chart (mean ± SEM) showing the area of AMPAR clusters within PSDs in control samples (Control: cluster area = 0.012 ± 0.001 $\mu m^2$). (**C**) Cumulative frequency distribution of nearest-neighbor distances (NNDs) between AMPAR-labeled immunogold particles in control samples (Control: n = 253 synapses; black) and samples from TLE patients (Epilepsy: n = 290 synapses; magenta). Cumulative frequency (%) is plotted as a function of NND (nm). Compared with the control distribution, the epilepsy distribution has a steeper slope, rather than being shifted leftward.
