## Supplemental Figure 3 for "Nanoscale remodeling of synaptic glutamate receptors and calcium channels in temporal lobe epilepsy"

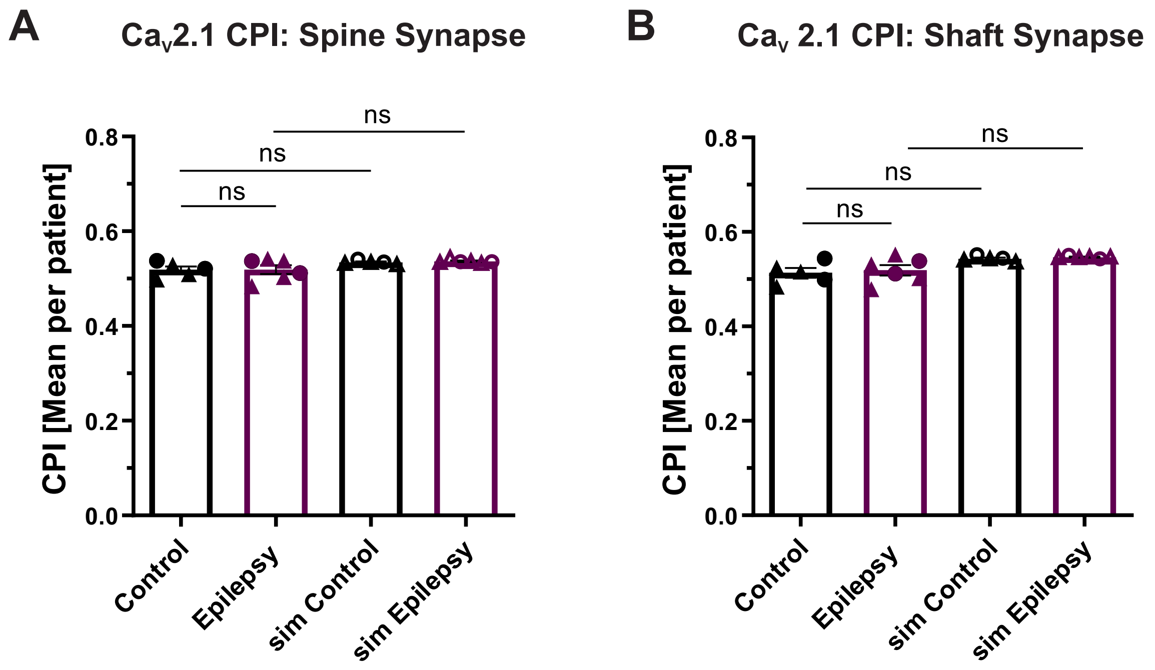


**Supplementary Figure 3: Distribution of $Ca_v$2.1 labeling within AZs of dendritic spine and shaft synapses in controls and TLE patients.** (**A**) Bar chart (mean ± SEM) of experimental Center-Periphery Index (CPI) of $Ca_v$2.1-labeled immunogold particles in spine synapses from controls (Control: CPI = 0.519 ± 0.007) and TLE patients (Epilepsy: CPI = 0.519 ± 0.010) indicating similar subsynaptic distribution patterns in both groups. It did not differ significantly from its corresponding simulated random distribution in controls (Control vs. sim Control: $P$ = 0.104) neither in TLE patients (Epilepsy vs. sim Epilepsy: $P$ = 0.108; two-way ANOVA with Holm-Šidák multiple-comparisons correction). (**B**) Bar chart (mean ± SEM) showing a similar subsynaptic distribution pattern in dendritic shaft synapses of controls (Control: CPI = 0.513 ± 0.010) and TLE patients (Epilepsy: CPI = 0.519 ± 0.011). No significant deviation from the respective simulated random distribution was observed in both controls (Control vs. sim Control: $P$ = 0.0634) and TLE patients (Epilepsy vs. sim Epilepsy: $P$ = 0.0634; two-way ANOVA with Holm-Šidák multiple-comparisons correction). ns, $P > 0.05$, not significant.
