## Supplemental Figure 4 for "Nanoscale remodeling of synaptic glutamate receptors and calcium channels in temporal lobe epilepsy"

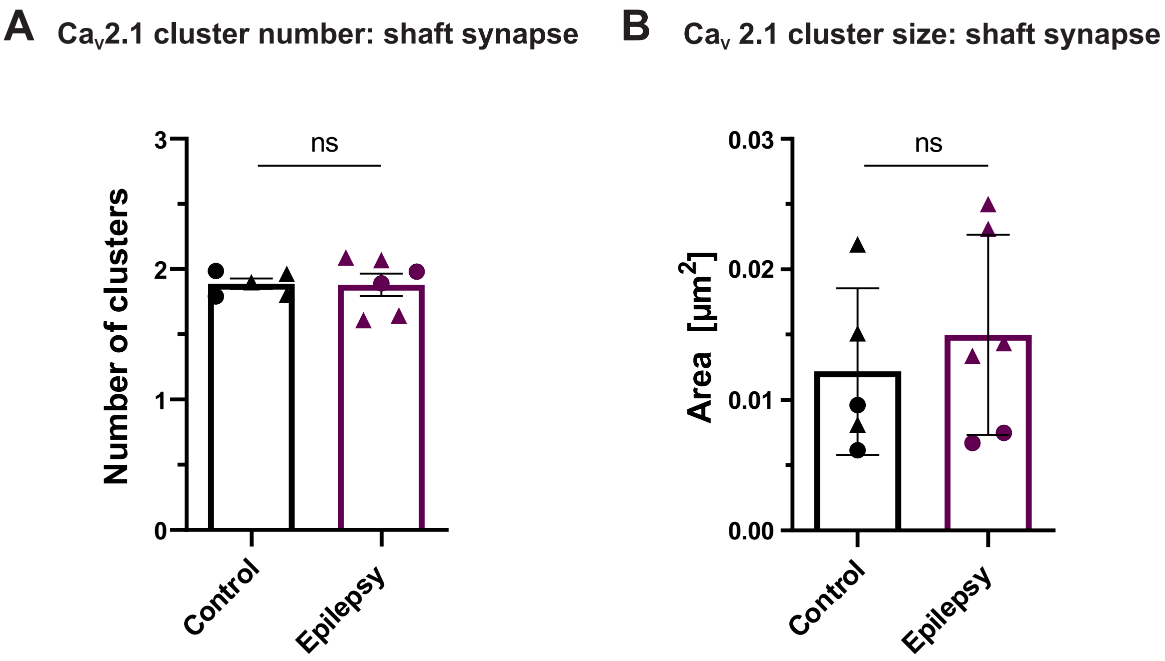


**Supplementary Figure 4: Clustering of $Ca_v2.1$ labeling within AZs of dendritic shaft synapses in controls and TLE patients.** (**A**) Bar chart showing the mean $Ca_v2.1$ cluster number within AZs of shaft synapses in controls (Control: n of clusters = 1.888 ± 0.040) and TLE patients (Epilepsy: n of clusters = 1.880 ± 0.086). No significant difference in the cluster number was observed ($P = 0.939$; unpaired t-test with Welch's correction). (**B**) Bar chart showing the mean $Ca_v2.1$ cluster area in controls (Control: cluster area = 0.012 ± 0.003 µm$^2$) and TLE patients (Epilepsy: cluster area = 0.015 ± 0.003 µm$^2$). No significant difference in the cluster area was observed ($P = 0.769$; unpaired t-test with Welch's correction). ns, $P > 0.05$, not significant.
